# Excitatory: inhibitory imbalance in Alzheimer’s disease is exacerbated by seizures with attenuation after rapamycin treatment in 5XFAD mice

**DOI:** 10.1101/2023.03.02.530499

**Authors:** AJ Barbour, S Gourmaud, E Lancaster, X Li, D Stewart, DJ Irwin, DM Talos, FE Jensen

## Abstract

Approximately 22% of Alzheimer’s disease (AD) patients suffer from seizures, and the co-occurrence of seizures and epileptiform activity exacerbate AD pathology and related cognitive deficits. Hence seizures may be a targetable component of AD progression. As epileptogenesis is associated with changes in neuronal excitatory: inhibitory (E:I) balance, we hypothesized that decreased markers of inhibition relative to those of excitation would be present in AD patients and exacerbated further when seizures were a comorbidity. We similarly hypothesized that an E:I imbalance would be present in five times familial AD (5XFAD) mice and augmented following pentylenetetrazol (PTZ) seizure kindling. AD temporal cortical tissue from patients with or without seizure history and brain tissue from 5XFAD mice were examined for changes in several markers of E:I balance, including the inhibitory GABA_A_ receptor, the chloride cotransporters, sodium potassium chloride cotransporter 1 (NKCC1) and potassium chloride cotransporter 2 (KCC2), and the excitatory NMDA and AMPA type glutamate receptors. We found that AD patients had decreased GABA_A_ receptor subunits and those with comorbid seizures had worsened cognitive and functional scores, and increased in NKCC1/KCC2 ratios, indicative of depolarizing GABA responses. The E:I imbalance appears to occur early in the disease course, as patch clamp recordings from CA1 neurons in hippocampal slices from prodromal 5XFAD mice showed decreased GABAergic inhibitory transmission and increased intrinsic excitability. In addition, seizure induction in prodromal 5XFAD mice further dysregulated NKCC1/KCC2, and altered the excitatory AMPA glutamate receptor protein expression, with a reduction in GluA2 subunit, indicative of calcium permeable-receptors. Finally, we found that chronic treatment with the mTORC1 inhibitor, rapamycin, at doses we have previously shown to attenuate seizure-induced α-amyloid pathology and cognitive deficits, could reverse aspects of E:I imbalance in these mice. Our data demonstrate novel mechanisms of interaction between AD and epilepsy and indicate that FDA-approved mTOR inhibitors hold therapeutic promise for AD patients with a seizure history.

## Introduction

Seizures can be a comorbidity in Alzheimer’s disease (AD) patients, with 10-22% showing clinically identifiable seizures and up to 64% displaying subclinical epileptiform activity^1^. Notably, the co-occurrence of seizures and epileptiform activity in AD patients enhances disease progression and worsens cognitive performance^1–5^. We and others have demonstrated bidirectional interactions with AD and epilepsy sharing similar pathology and convergence upon common underlying cellular mechanisms. Human temporal lobe epilepsy brain tissue shows increased Aβ plaques and increased hyperphosphorylated tau (pTau)^6–8^ and the accumulation of these neuropathological proteins in AD animal models are associated with elevated neuronal hyperexcitability and seizure susceptibility ^9–14^. Further, we recently demonstrated that a history of seizures is associated with enhanced Aα and pTau pathology in human AD^14^.

While mechanisms of seizure generation in AD are not fully elucidated, research points to evidence of a synaptic excitatory:inhibitory (E:I) imbalance in chronic epilepsy, including altered expression and function of certain neurotransmitter receptors and ion cotransporters^15–17^. Chronic epilepsy human and animal model studies show evidence of reduced inhibitory neurotransmission, with decreases in the ratio of α1/α3 inhibitory gamma-aminobutyric acid type A receptor (GABA_A_R) subunits that diminish inhibitory postsynaptic currents (IPSCs)^18–21^. Inhibitory GABAergic transmission is further attenuated by paradoxical depolarizing GABA_A_R currents due to reversal of the chloride (Cl^-^) gradient caused by increases in the ratio of the sodium (Na^+^) potassium (K^+^) Cl^-^ co-transporter 1 (NKCC1), which imports Cl^-^, to that of the potassium Cl^-^ cotransporter 2 (KCC2), which exports Cl^-22–25^. Commensurate with decreased inhibitory drive in epilepsy, there is an increase in glutamate-receptor mediated excitability, in part due to alterations in α-amino-3-hydroxy-5-methyl-4-isoxazolepropionic acid receptor (AMPAR) and N-methyl-D-aspartate receptor (NMDAR) subunits expression, with increases in the ratios of GluA1/GluA2 AMPAR subunits and GluN2B/GluN2A NMDAR subunits resulting in GluA2-lacking, Ca^2+^-permeable AMPAR^26–28^ and extended decay times and elevated Ca^2+^ influx, respectively^29–31^.

Pathologic E:I alterations are not unique to epilepsy, as evidence for neuronal hyperexcitability has been seen in other neurodevelopmental and neurodegenerative disorders^32–35^. In AD patients, altered GABA_A_R subunit regulation has been observed^36^, including decreased α1/α3 expression^37^. Further, evidence for paradoxically depolarizing GABA currents have been found, with reduced KCC2 in the AD11 mouse model^38^ and elevations of NKCC1 after Aα_1-42_ hippocampal injection^39^. Aα_1-42_ has also been shown to increase Ca^2+^-permeable AMPARs which can result in memory deficits via reduced long-term potentiation and synapse loss^40–43^. However, given the prevalence of network excitability and seizures in AD, and the evidence that seizures can alter E:I balance in epilepsy, understanding the impact of seizures and the evolution of neuronal hyperexcitability in AD progression may lead to new mitigatable therapeutic E:I targets. To understand the progression of E:I balance, we examined the 5XFAD mouse model at two stages. We also examined E:I balance in human AD with known seizure history and after seizure induction in 5XFAD mice to determine the impact of seizures on E:I balance in AD.

Preclinical and clinical epilepsy research has revealed several candidate signaling pathways that might link E:I imbalance and network hyperexcitability in epilepsy to neurodegeneration. One example is the substantial evidence for the activation of the mammalian target of rapamycin complex 1 (mTORC1) as a key common element between epilepsy and neurodegeneration. mTORC1 mediates pathological tau and APP processing^44–48^ and can regulate excitatory and inhibitory receptor and network function^49,50^. Studies of models of Tuberous Sclerosis Complex, characterized by constitutive mTORC1 overactivation commonly resulting in epilepsy^51,52^, have demonstrated that blockade of mTORC1 with rapamycin can rescue GABAergic neurotransmission and epilepsy^53–56^. Rapamycin treatment has also rescued neuronal hyperexcitability in models of autism spectrum disorder^57^ and neuropathic pain^58^. Together, these results indicate that targeting mTORC1 activity might restore E:I balance across multiple disease models, including AD.

Considering these data and that rapamycin and other rapalogs are approved by the United States Food and Drug Administration for other indications, mTORC1 is a promising target for therapeutic intervention for patients with AD and co-morbid epilepsy. Indeed, we recently found that AD patients with a history of seizure display enhanced tau and Aα pathology, associated with increased mTORC1 activity^14^. In addition, we observed that the induction of seizures in the 5XFAD mice exacerbates Aα pathology and cognitive deficits, and that these changes are reversible with chronic low-dose rapamycin treatment.

Given that E:I imbalance may contribute to the cognitive impairments observed in AD, we hypothesized that key components of E:I balance may be dysregulated in AD and that these alterations emerge early in the disease course and may be exacerbated by seizures. First, we measured established markers of E:I imbalance in largely the same human control and AD temporal neocortex from patients with and without known seizure history as our previous study^14^ and indeed found greater alterations in several components of the GABAergic system, along with worsened cognitive and functional performance in AD patients with a seizure history. Next, we examined the 5XFAD mouse model for protein and electrophysiological indications of hyperexcitability at the prodromal stage (∼4 months) and directly assessed the effects of seizures induced at this stage on E:I markers at a later, chronic timepoint (7 months of age). Finally, given that we found protective effects on pathology and cognition with rapamycin in this model^14^, we utilized the tissue from our prior study to examine whether these therapeutic benefits extend to markers of E:I balance. Overall, we found that E:I imbalance began at a prodromal stage in 5XFAD mice and that seizures significantly exacerbated this imbalance, while chronic rapamycin treatment significantly attenuated these effects.

## Materials and Methods

### Human subjects

All protocols and procedures were approved under the ethical standards of the Institutional Review Board of the University of Pennsylvania (Philadelphia, PA). The AD cognitive study population consisted of 105 patients, selected by querying the integrated neurodegenerative disease database^59^ of the Center for Neurodegenerative Disease and Research (CNDR) at the University of Pennsylvania (Philadelphia, PA) for available global clinical dementia rating (CDR) score and CDR Sum of Boxes (CDR-SOB) closest to death. All patients were enrolled in observational research at Penn, with standardized assessments and medical history that includes assessment of seizure history ^60^. Of the AD cases, 18 (6 males and 12 females; mean age at death= 77.38 years) had a reported clinical seizure history (AD+Sz). The remaining 87 AD patients (43 males and 44 females; mean age at death= 79.7 years) had no known seizure history (AD-Sz). AD clinical diagnosis was established during life based on the clinical history, neurological and neuropsychological assessment, and confirmed by post-mortem histopathological staging of AD neuropathological markers (Aβ_42_ and tau pathology), according to the National Institute on Aging-Alzheimer’s Association (NIA-AA) guidelines^60^. Detailed methods for neurocognitive assessments can be found in Supplementary Materials.

For a subset of the population detailed above, post-mortem brain specimens were acquired from CNDR and consisted of superior/mid-temporal cortex frozen samples (n= 34) and formalin-fixed paraffin-embedded sections (n=15). Detailed clinical characteristics of this AD cohort, obtained from the CNDR INDD database, were previously published^14^ and included age at onset of cognitive decline (disease duration), cause of death, brain weight and ordinal rating of gross ventricular enlargement at the time of death, Braak and Thal stages, seizure history and medications. Of the AD cases, 14 (6 males and 8 females; mean age at death= 78.4 years; mean PMI= 11h) had a reported clinical seizure history (AD+Sz). The other 20 AD patients (10 males and 10 females; mean age at death= 73.9 years; mean PMI= 12h) had no known seizure history (AD-Sz). Region-matched control tissue (n=15; 11 males and 4 females; mean age at death= 62.4 years; mean PMI=15.8 hours), received from the CNDR and NIH NeurobioBank, had no known neurologic or psychiatric history. AD patients were significantly older than controls at time of death (p<0.001). PMI did not differ between control and AD groups. Clinical characteristics for all patients included in the cognitive and biochemical studies can be found in Table S1.

### Mice

All animal procedures were approved and performed in accordance with the guidelines of the Institutional Animal Care and Use Committee (IACUC) Office of Animal Welfare of the University of Pennsylvania (Philadelphia, PA). Male and female 5XFAD and WT mice on a mixed B6SJL background second generation (F2) generated from mice originally purchased from Jackson Laboratory (MMRC stock #3840, Bar Harbor ME) were used in these studies. 5XFAD mice harbor the human APP and PSEN1 genes under the Thy1 promoter containing five mutations for aggressive amyloid development (APP: Swedish K670N/M671L, Florida I7516V, and London V717I; PSEN1: M146L and L286V). Genotyping was conducted as specified by Jackson Laboratory. Mice were randomly allocated to treatment groups, which were counterbalanced for sex and litter. No mice were excluded from analysis.

### PTZ kindling

PTZ kindling was performed as previously described ^14^. Briefly, mice were administered eight sub-convulsive doses of the GABA_A_R antagonist PTZ (i.p., 35 mg/kg of PTZ, Sigma-Aldrich, St Louis, MO) or vehicle (NaCl 0.9%, Sigma-Aldrich) every 48 h for 15 days. Mice were monitored and video-recorded for one hour after PTZ administration and scored for seizure severity to confirm kindling^14^. One month post-kindling, mice were administered the same dose of PTZ (35 mg/kg) or vehicle to confirm the development of kindling. In our therapeutic cohort, we used a slightly modified kindling procedure to minimize animal loss. Mice were removed from kindling when they reached tonic-clonic seizures on three consecutive days of treatment and were not administered PTZ one month post kindling.

### Rapamycin treatment

Mice were administered encapsulated rapamycin feed dosed at 14mg/kg (5LG6 w/151ppm Encapsulated rapamycin Irr, Emtora Holdings, San Antonio, TX) or control chow (5LG6 w/151ppm Eudragit Irr, Emtora Holdings)^14,61–63^ beginning at completion of PTZ kindling (3.5 months of age) until euthanasia at seven months of age. Mice were treated with rapamycin 2.24mg/kg/day by ad libitum consumption of 167 mg/one gram body weight/day.

### Electrophysiology

Mice, 4-5 months of age, were anesthetized with sodium pentobarbital (90 mg/kg, i.p.) and perfused with ice-cold, oxygenated, artificial cerebrospinal fluid (aCSF, in mM, 124 NaCl, 2.5 KCl, 1.2 NaH_2_PO_4_, 24 NaHCO_3_, 5 HEPES, 2 CaCl2. 2 MgSO_4_, and 12.5 glucose, pH 7.3-7.4). The brains were removed and cut horizontally (300 µm) in ice-cold, oxygenated, NMDG-aCSF solution (in mM, 93 N-methyl-D-glucamine (NMDG), 2.5 KCl. 1.2 NaH_2_PO_4_, 30 NaHCO_3_, 20 HEPES, 0.5 CaCl_2_, 10 MgSO_4_, 25 glucose, 2 thiourea, 5 sodium ascorbate, and 3 sodium pyruvate, pH 7.3-7.4) using a Leica VT1000s vibratome. Slices were kept in an NMDG-aCSF at 34 °C for 30 min. During this time, incremental amounts of 2M NaCl were added to gradually increase the Na^+^ concentration in the NMDG-aCSF ^64^.Then, slices were transferred to aCSF at room temperature and kept until used for recording.

Whole-cell patch clamp recording was performed from hippocampal CA1 pyramidal neurons, identified by a Zeiss Axioscope 2 equipped with IR-DIC. Slices were superfused with room temperature aCSF at a rate of ∼ 2ml/min. Signals were amplified and digitized with an Axopatch 200B and a Digidata 1440A. Data were collected and analyzed using pCLAMP 10, Clampfit 10.2 (Molecular Devices) and Mini Analysis 6.0 (Bluecell). For testing action potential (AP) firing properties in current clamp mode, patch pipette electrodes (2-3 MΩ) were filled with internal solution composed of, in mM: 126 K gluconate, 4 KCl, 10 HEPES, 0.3 EGTA, 0.3 Na-GTP, 3 Mg-ATP, 10 phosphocreatine, pH 7.3, Osm 290. For recording miniature inhibitory synaptic currents (mIPSCs), cells were voltage clamped at −60 mV using electrodes filled with internal solution composed of, in mM; 130 CsCl, 4 NaCl, 10 HEPES, 0.3 EGTA, 1 MgCl2, 0.3 Na-GTP, 3 Mg-ATP, 10 phosphocreatine, pH 7.3, Osm 290. To isolate mIPSCs, TTX (1 µM), NBQX (20 µM), and AP-5 (50 µM) were added to superfusing aCSF. Cells with membrane resistance above 100 MΩ and series resistance less than 10 MΩ were accepted for analysis.

### Western Blot

Mice were anesthetized with 50 mg/kg pentobarbital (Sagent Pharmaceuticals, Schaumburg, IL) and perfused intracardially with cold PBS. Hippocampi and cortices were dissected and immediately frozen. Both human and mouse brain samples were homogenized in a lysis buffer containing 1.1% sucrose, 50 mM Tris HCl pH 7.5, 500 μM CaCl_2_, 1 mM MgCl_2_, 1 mM NaHCO_3_, 1X protease inhibitor, 1mM phenylmethanesulfonyl fluoride (PMSF), and 1X HALT^TM^ (Thermo Fisher Scientific, Carlsbad, CA) on ice and centrifuged at 4,100 g for 10 min at 4° C. Membrane fractions were generated by further centrifugation at 3,000 g for 10 min at 4° C. The resultant supernatant was centrifuged at 13,000 g for 10 min at 4° C. The pellet was reconstituted in lysis buffer. Protein concentrations were determined using the Bradford Protein Assay kit (Bio-Rad, Hercules, CA). Proteins were separated by electrophoresis on 4-20% SDS-PAGE gels (Bio-Rad) and transferred to polyvinylidene difluoride membranes (EMD Millipore, Burlington, MA) following previously used standard protocols^6,14^. Primary antibodies are listed in Table S2. Detected proteins were imaged with Odyssey Imaging System (LI-COR Biosciences) and quantified with Image Lab (Bio-Rad). Each protein was normalized to β-actin expression level.

### Statistics

For cognitive and functional performances, comparisons between AD+Sz and AD-Sz for ranked ordinal data were made by Mann-Whitney test. For the human tissue studies, all AD to control comparisons were made by unpaired t-tests or Mann-Whitney tests for non-normal data. Control, AD-Sz, and AD+Sz comparisons were made by one-way analysis of variance (ANOVA) or Kruskal-Wallis for non-normal data. For the prodromal mouse study, 5XFAD to WT comparisons were made by unpaired t-test. For the nontherapeutic PTZ study, comparisons were made by two-way ANOVA with genotype and kindling as independent variables. For the therapeutic PTZ study, 5XFAD and WT mice were analyzed separately by two-way ANOVAs with PTZ and rapamycin treatment as independent variables. All ANOVA analyses were followed by Tukey’s post hoc test to examine group differences when main effects or interactions were found. Additional multiple linear regressions were performed for all data to determine potential sex effects. Where sex effects were present, data points are displayed as squares (female) or triangles (male) to visualize these differences. Correlations were calculated by simple linear regression. We considered results to be significant at p<0.05. All statistical analyses were performed in GraphPad Prism 9 software (San Diego, CA).

### Data Availability

The data that support the findings of this study are available from the corresponding author upon reasonable request.

## Results

### AD patients with seizure history have worsened cognitive and functional performance

To establish potential associations between seizures and cognitive and functional deficits in AD, we examined scores from clinical (global CDR and CDR-SOB) and caretaker questionnaire (DSRS) for subjects with known seizure history. The AD+Sz group showed elevated final global CDR (p<0.001, n=18-87, Figure 1A) and CDR SOB (p<0.0001, n=18-87, Figure 1B) compared to the AD-Sz group. AD+Sz patients also showed significantly worsened overall functional rating (DSRS) compared to those in the AD-Sz group (p<0.01, n=14-19, Figure 1C), including significantly worse ratings in memory, recognition of family members, mobility, orientation to place, social and community activities, and incontinence (p<0.05, n=14-19, Figure S1). These data indicate seizures are associated with worsened cognitive performance and overall function, corroborating previous studies examining epileptiform activity in AD^2–5^ and suggesting heightened neuronal dysfunction.

**Figure 1.**
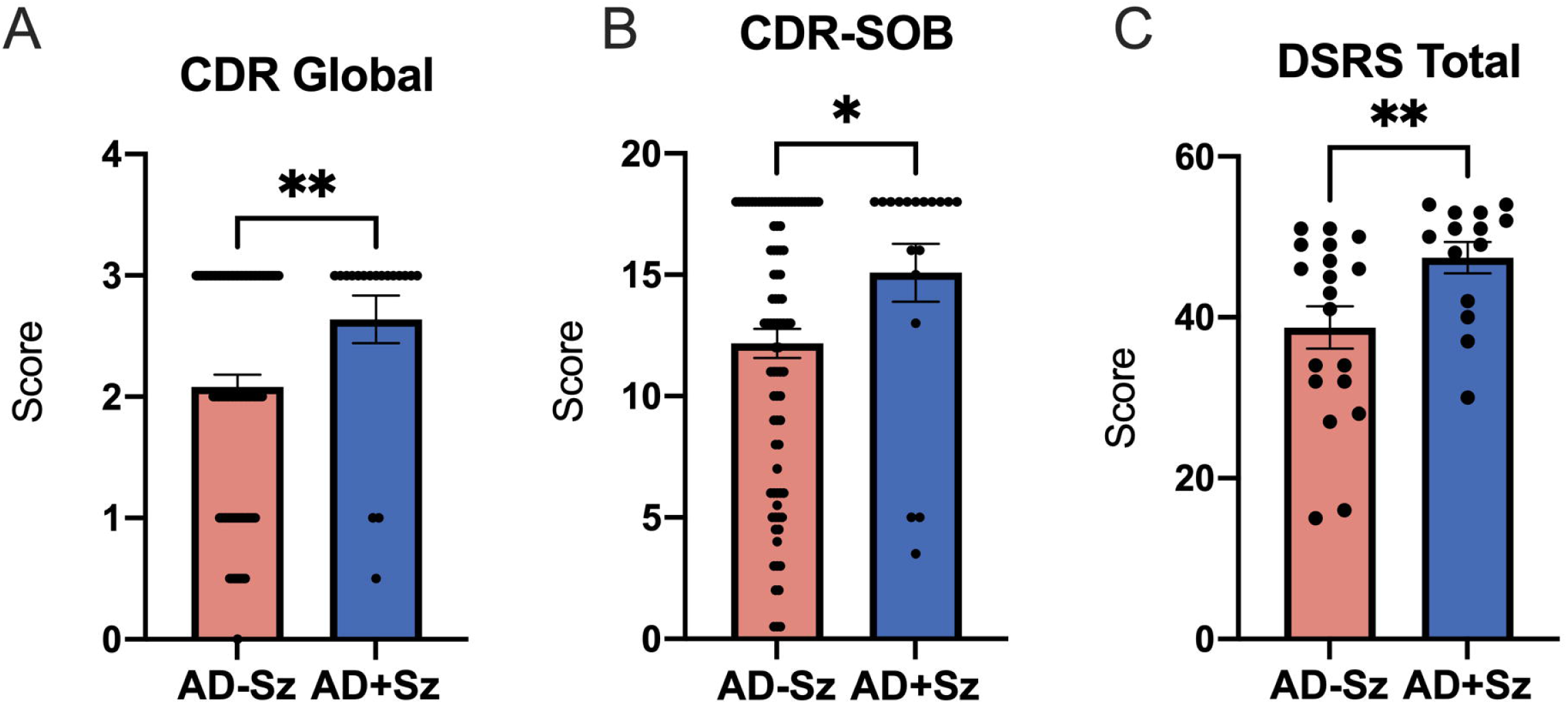
Seizure history is associated worse cognitive and functional performance in AD. Comparisons between AD-Sz and AD+Sz in Final global CDR (**A**), CDR sum of boxes (**B**), and DSRS Total (**C**). CDR: n=18 (AD+Sz) – 87 (AD-Sz); DSRS: n=14 (AD+Sz) – 19 (AD-Sz). *p<0.05, **p<0.01.

### AD patients with a seizure history exhibit more severe dysregulation of proteins that mediate excitation: inhibition balance

We hypothesized that the worsened functional and cognitive deficits found in AD+Sz patients would be associated with worsening of markers of E:I imbalance. Thus, we examined ratios of α1/α3 subunits of the inhibitory GABA_A_R^19^ and Cl^-^ cotransporters NKCC1/KCC2^22,23,65^. We found a significant decrease in GABA_A_Rα1 expression in AD+Sz compared to controls (F=3.52, Tukey’s post hoc: p<0.01, Figure 2B), that was not found in AD-Sz, and which drove the reduction in GABA_A_Rα1 found across all AD patients (p<0.05, Figure 1B). This alteration also resulted in significantly decreased GABA_A_Rα1/α3 ratio in all AD patients when compared to controls (p<0.01, Figure 1B). The ratio of NKCC1/KCC2 was also significantly increased in the AD+Sz group when compared to the AD-Sz group (p<0.05, Figure 2C). This difference was largely attributable to downregulation of KCC2 in the AD+Sz group, compared to the AD-Sz group (F=3.3, Tukey’s post hoc: p<0.05, Figure 2C). In addition, sex was a significant predictor of KCC2 dysregulation, with decreased levels in males (p<0.05, [0.0314, 0.230], Figure 2C). Taken together these results suggest AD+Sz patients had greater GABAergic dysfunction due to reduced hyperpolarization and indications of depolarizing GABA_A_R-mediated signals.

**Figure 2.**
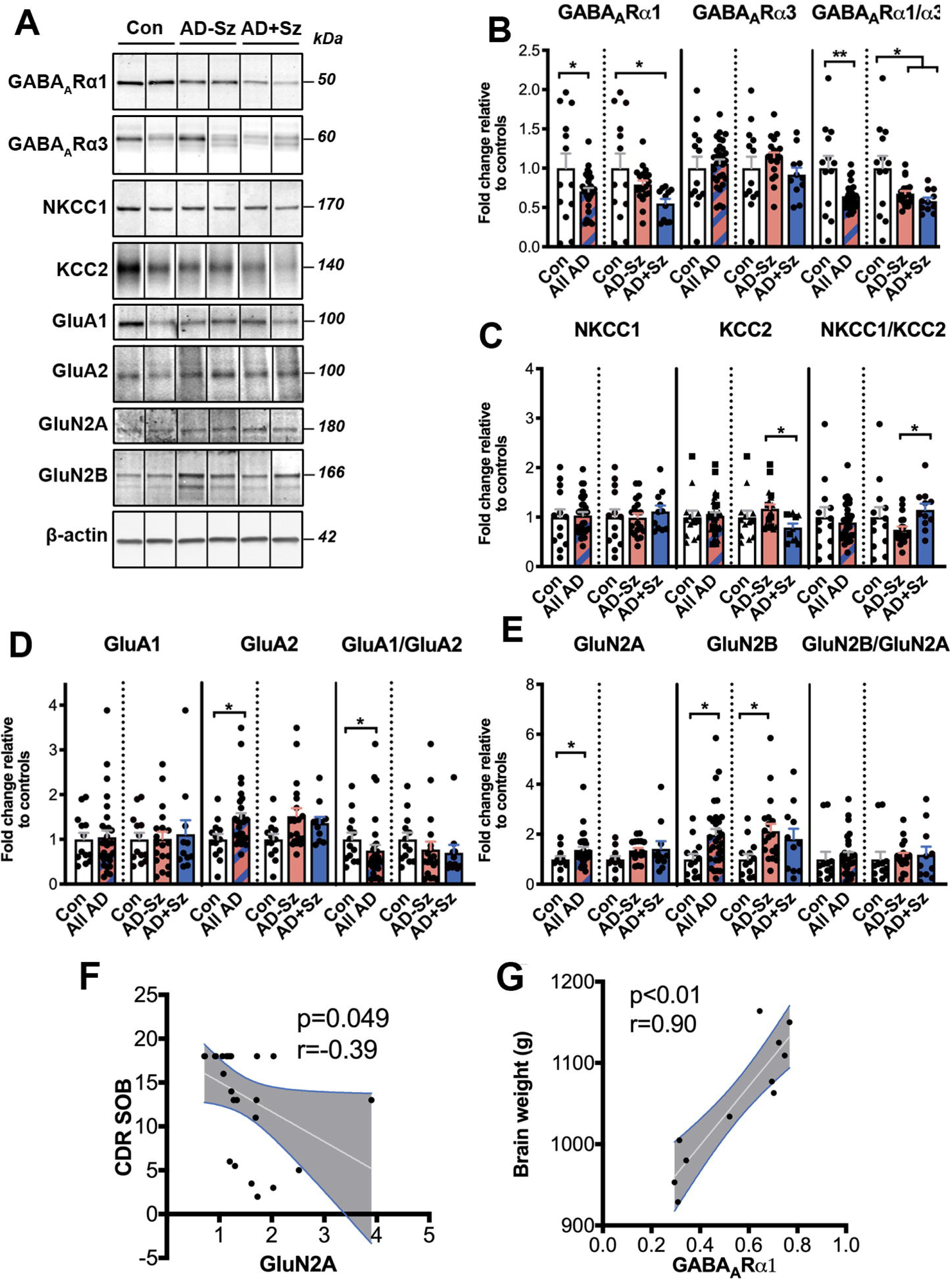
Dysregulation of proteins involved in excitatory: inhibitory balance in AD patients with and without seizure history. (**A**) Representative western blot images for (B-E) showing non-adjacent bands originating from the same blot of the temporal cortex from control cases (Con; n=13) and Alzheimer’s disease patients (AD; n=30) split into subgroups, those without (AD-Sz; n=19) or with (AD+Sz; n=11) known seizure history. Semi-quantitative analysis of (**B**) GABA_A_R subunits GABA_A_Rα1 and GABA_A_Rα3 and corresponding ratio GABA_A_Rα1/GABA_A_Rα3; (**C**) Cl^-^ cotransporters NKCC1 and KCC2 and corresponding ratio NKCC1/KCC2; (**D**) AMPAR subunits GluA1 and GluA2 and corresponding ratio GluA1/GluA2; (**E**) NMDAR subunits GluN2A and GluN2B and corresponding ratio GluN2B/GluN2A. *p<0.05, **p<0.01. (**F**) Pearson correlations in all AD showing relationship between GluN2A and CDR SOB score and (**G**) showing the relationship between GABA_A_Rα1 expression and brain weight at death in grams in AD+Sz patients. Grey areas indicate 95% confidence interval for the two means.

To further explore inhibitory dysfunction, we compared levels of the predominant PV-expressing interneurons^66^ in tissue from AD+Sz and AD-Sz, given that there is a well-established association between PV cell loss, cognition^67^ and epilepsy^68,69^. Immunohistochemistry revealed that AD+Sz had significantly decreased PV+ cells compared to controls (p<0.01, Figure S2 A-B) that was not found in AD-Sz which drove the significant decrease in PV+ cells found between all AD and controls (p<0.01, Figure S2 A-B).

To determine if neuronal excitation is heightened in AD and whether seizures cause further alterations, we examined the expression of AMPA and NMDA glutamate receptor subunit proteins, namely ratios of GluA1/GluA2 and GluN2B/GluN2A subunits^26,27,29^ (Figure 2A-E). While no differences were found between AD+Sz and AD-Sz in AMPAR subunits or subunit ratios (Figure 2D), we found increased expression of GluA2 subunits in AD temporal cortex overall compared to controls (p<0.05, Figure 2D) and the corresponding GluA1/GluA2 ratio was decreased in the AD cases compared to controls (p<0.05, Figure 2D). In examining NMDAR subunits, there was no difference in GluN2B or GluN2A levels nor in the ratio of GluN2B/GluN2A between the AD+Sz and AD-Sz groups (Figure 2E). However, GluN2A and GluN2B subunits were both expressed at significantly higher levels in AD temporal cortex when compared to controls (p<0.05 and p<0.001, respectively, Figure 2E).

Finally, we performed linear regressions between clinical outcomes of AD and markers of E:I balance. We found that CDR SOB was inversely correlated with GluN2A levels across all AD subjects (r=-0.39, p<0.05, Figure 2F) while gross brain atrophy ratings at autopsy was associated with decreased levels of GABA_A_Rα1 in AD+Sz (r= 0.90, p<0.01, Figure 2G), indicating an interaction between these markers of E:I balance and cognitive decline and AD severity, respectively. No correlations were found between markers of E:I balance and pathology previously measured in these samples^14^. These results provide strong evidence of decreased inhibition and increased excitation in AD patients overall, regardless of seizure history, and an exacerbation of selective regulators to increase GABAergic dysfunction in the AD+Sz group.

### Excitation: inhibition imbalance in prodromal stage 5XFAD mice

Given that our human tissue samples are limited by the fact they are from patients at late-stage AD (Table S1), an animal model is ideal to examine earlier stages. Indeed, there is evidence of E:I imbalance reported in several AD mouse models at early stages^70–72^. We thus used the 5XFAD mouse model to determine whether baseline E:I imbalance is present at the prodromal stage (4 months of age), when mild behavioral impairment^73^, AD pathology^74^, subclinical epileptiform electroencephalographic activity^75^, and decreased thresholds for chemoconvulsant induced seizures^14^ are present.

Western blot revealed early alterations in markers for GABA inhibition, with 5XFAD mice exhibiting decreased GABA_A_Rα1/GABA_A_Rα2 ratio in the hippocampus (p<0.05, n=14-15), and a trend towards decreased KCC2 in the cortex (p=0.09, n=14-15). Notably, no changes in excitatory glutamate AMPAR and NMDAR subunits in hippocampus and cortex remained unchanged at this prodromal stage (Figure 3).

**Figure 3.**
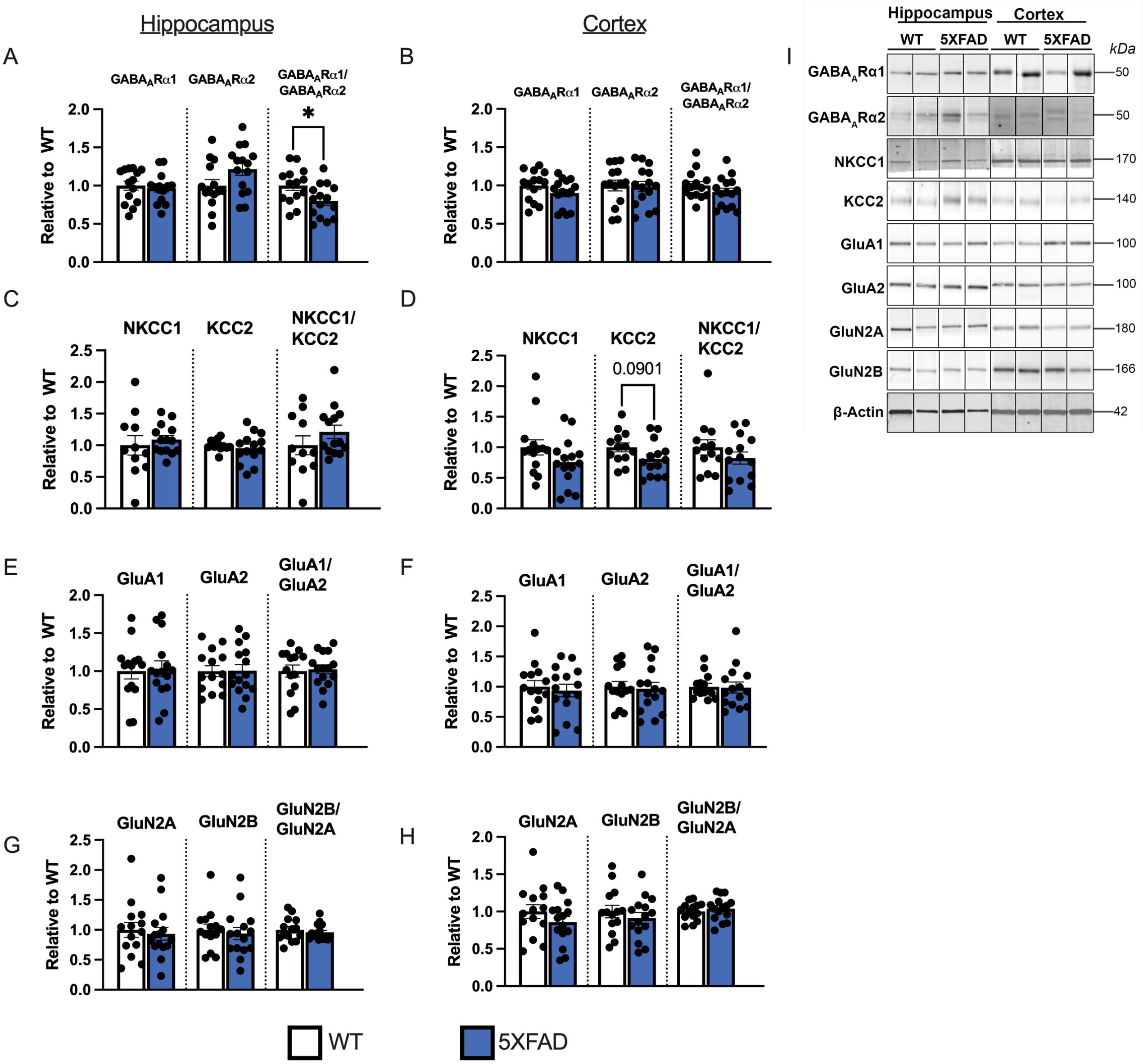
Excitatory: inhibitory markers in prodromal 5XFAD mice. Analysis of bands from western blots for E:I markers from the hippocampus (A, C, E, G) and cortex (B, D, F, H) of 5XFAD and WT mice. (**A, B**) GABA_A_R subunits GABA_A_Rα1 and GABA_A_Rα2 and corresponding ratio GABA_A_Rα1/GABA_A_Rα2 (**C, D**) Cl^-^ cotransporters NKCC1 and KCC2 and corresponding NKCC1/KCC2 ratio; (**E, F**) AMPAR subunits GluA1 and GluA2 and corresponding GluA1/GluA2 ratio; (**G, H**) NMDAR subunits GluN2A, GluN2B, and corresponding GluN2B/GluN2A ratio. (**I**) Representative Western blot images for (**A-H**) showing non-adjacent bands originating from the same blot. Error bars represent standard error of the mean. n=11-15 per group. *p<0.05.

Given that decreases in GABA_A_Rα1/α2 ratio has been reported to decrease inhibitory synaptic activity in epilepsy models^18–21^, we next aimed to determine whether these protein changes also resulted in similar deficits in inhibitory synaptic activity the prodromal 5XFAD mice. We thus measured mIPSCs using whole-cell patch clamp in CA1 pyramidal neurons in hippocampal slices from 4 month old prodromal 5XFAD mice. Indeed, mIPSC amplitudes were significantly decreased in 5XFAD compared to WT mice (p<0.0001, number of mIPSC events=8316 and 8086, n=16 cells/groups, 2 mice/group, Figure 4A, B, C), consistent with reduced GABA_A_Rα1/GABA_A_Rα2 ratio^21^. Importantly, there was no change in mIPSC frequency, confirming postsynaptic mechanisms for inhibitory dysfunction. These changes in GABAR function were also associated with increased network excitability in 5XFAD mice as measured by recording evoked action potentials (APs) in current clamp mode using depolarizing step currents of 800 ms duration in the aforementioned CA1 pyramidal neurons (Figure S3). Overall, these data confirmed that the altered GABA expression in hippocampi of prodromal 5XFAD mice was associated with hyperexcitability and may underlie increased seizure susceptibility that we have previously found at this stage in 5XFAD mice^14^.

**Figure 4.**
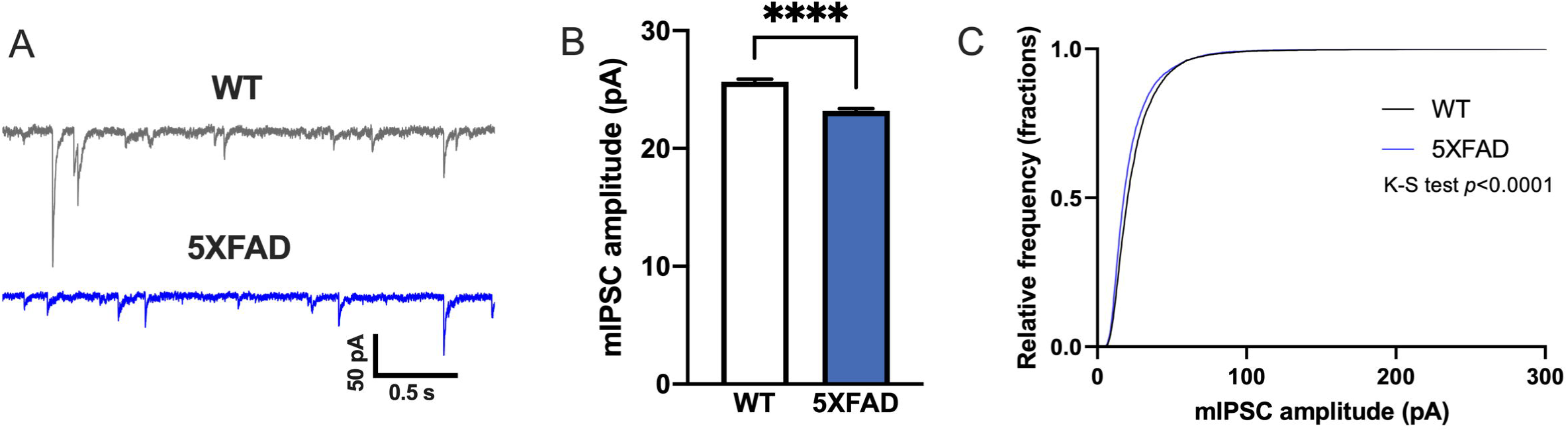
Inhibitory dysfunction in pyramidal neurons from prodromal 5XFAD CA1. (**A**) Representative mIPSC traces from WT and 5XFAD. (**B**, **C**) Quantifications of mIPSC amplitudes. n= 16 cells from 2 WT mice, 16 cells from 2 5XFAD mice. ****p<0.0001.

### Kindling during the prodromal stage alters the progressive age-related dysregulation of proteins involved in excitation: inhibition balance in 5XFAD mice

Given the findings of enhanced E:I imbalance in human tissue from AD patients with seizures, we sought to determine how E:I balance changes with disease progression in 5XFAD mice, and whether chronic seizures are associated with further dysregulation of E:I imbalance. We performed PTZ seizure kindling or control protocols in prodromal (3.5 month old) 5XFAD and littermate WT mice and analyzed expression levels of markers of E:I imbalance in the 7 month old (3.5 months post kindling) hippocampus (Figure 5) and cortex (Figure S4). 5XFAD mice had decreased hippocampal GABA_A_Rα1/α3 ratio compared to WT mice (genotype effect: F_1,61_ = 22.76, p<0.0001, Figure 5A), significant decreases in both non-kindled (p<0.05) and PTZ-kindled 5XFAD mice (p<0.01). No differences in GABA_A_Rα1/α3 were found due to kindling. The changes in hippocampal GABA_A_R ratio seem to be largely driven by decreases in the GABA_A_Rα1 subunit in 5XFAD mice (genotype effect: F_1,61_=40.09, p<0.0001) (Figure 5A). Decreased GABA_A_Rα1/α3 ratio were also found in the cortex (interaction: F_1,43_ = 4.25, p<0.05), in non-kindled 5XFAD mice compared to WT (p<0.05) (Figure S4A). These changes were similar to patterns seen in human AD brain (Figure 2) and prodromal (4 month) 5XFAD tissue (Figure 4). In addition, sex was a significant predictor of hippocampal GABA_A_Rα1/α3 with greater decreases found in females (p<0.05, [0.0156, −0.1591], Figure 5A). Taken together, these data demonstrate persistent GABA_A_R alterations into later disease stages in 5XFAD mice and may contribute to continued heightened seizure susceptibility previously demonstrated in these 5XFAD mice^14^.

**Figure 5.**
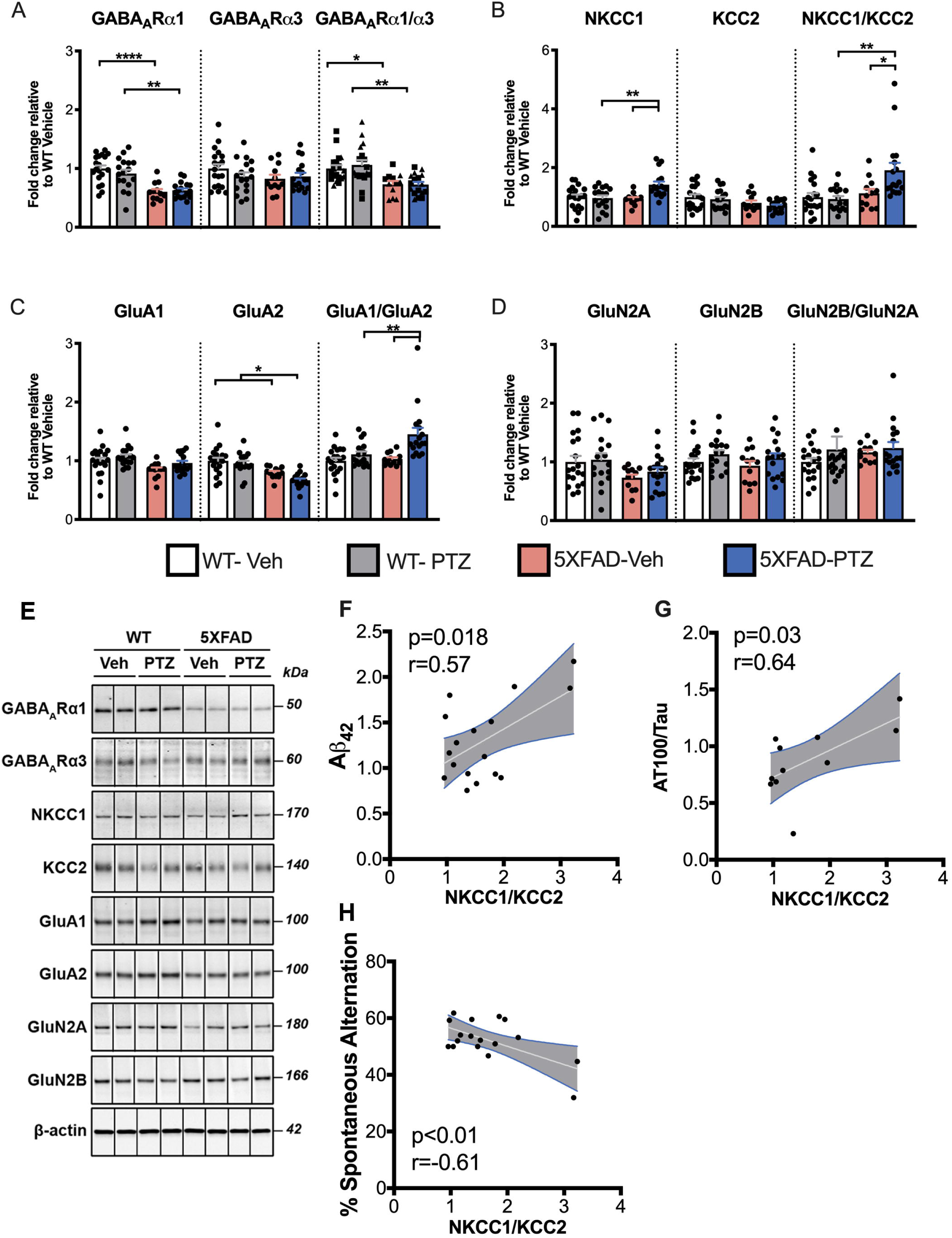
Excitatory:inhibitory imbalance in the hippocampus of 5XFAD mice following induced seizures. (**A-E**) Quantification of (**A**) GABA_A_R subunits GABA_A_Rα1 and GABA_A_Rα3 and corresponding ratio GABA_A_Rα1/GABA_A_Rα3; (**B**) Cl^-^cotransporters NKCC1 and KCC2 and corresponding ratio NKCC1/KCC2; (**C**) AMPAR subunits GluA1 and GluA2 and corresponding ratio GluA2/GluA1; (**D**) NMDAR subunits GluN2A and GluN2B and corresponding ratio GluN2B/GluN2A. (**E**) Representative western blot images for (A-D) showing non-adjacent bands originating from the same blot. (**I**) Pearson correlation of PTZ kindled 5XFAD mice showing the relationship between NKCC1/KCC2 and Aβ normalized to 5XFAD-Veh, (**F**) pTau AT100(Thr212, Ser14):Tau normalized to WT-Veh (**G**), and % spontaneous alternations in the Y-maze. Group comparisons for Aβ ELISA, pTau western blots and the Y-maze were previously published ^14^ (**H**). Grey areas indicate 95% confidence interval for the two means. n = 12-19 WT-vehicle, 12-17 WT-PTZ, 12 5XFAD-vehicle and 17 5XFAD-PTZ. *p<0.05, **p<0.01. Females and males are designated by square and triangle data points, respectively, where sex effects were found.

We next tested whether there were age-related and/or kindling induced alterations to Cl^-^ cotransporters in 5XFAD mice. As opposed to the prodromal stage, 7-month-old 5XFAD mice showed a decrease in hippocampal expression of the Cl^-^ exporter, KCC2, regardless of PTZ treatment (genotype effect, F_1,61_ = 12.35, p<0.001), suggesting worsened inhibitory deficits commensurate with pathological progression. In addition, kindling resulted in an increased NKCC1/KCC2 ratio in the hippocampus of 5XFAD mice (interaction: F_1,61_=5.78, p<0.05) compared to WT and non-kindled 5XFAD mice (p<0.05. Figure 5B), due to a significant increase in expression of the Cl^-^ importer, NKCC1 (interaction: F_1,61_=7.93, p<0.01; Tukey’s post hoc: p<0.01) (Figure 5C). In the cortex (Figure S4), we also found a significant increase in NKCC1/KCC2 ratios due to PTZ kindling (kindling effect: F_1,43_=4.49, p<0.05), which was largely driven by a reduction of KCC2 in kindled WT mice (interaction: F_1,43_= 6.75, p<0.05). Interestingly, there was a decrease in NKCC1 in the non-kindled 5XFAD cortex (genotype effect: F_1,43_= 5.2, p<0.05) compared to non-kindled WT mice (Tukey’s post hoc: p<0.05), which may indicate a compensatory mechanism, although NKCC1/KCC2 ratios were not significantly different (Figure S3B).These data suggest that depolarizing GABA^76^ may contribute to the exacerbation of GABAergic dysfunction in kindled 5XFAD mice.

We next sought to determine whether glutamate receptors may be dysregulated as a function of disease progression and also by kindled seizures in 5XFAD mice. In the hippocampus, we found a significant effect to increase GluA1/GluA2 ratios in PTZ kindled hippocampus (genotype × kindling, F_1,61_=4.622, p<0.05) (Figure 5C, p<0.05). Increased GluA1/GluA2 ratio is a source of E:I imbalance that has been described in both human cases and mouse models of status epilepticus^29^. In addition, both subunits were decreased in the hippocampus of 5XFAD mice regardless of kindling status (GluA1 genotype effect, F_1,61_ = 7.722, p<0.01; GluA2 genotype effect, F_1,61_ = 30.28, p<0.0001). Similar alterations in AMPAR subunit expression and ratios were found in the cortex (Figure S4C). The NMDAR GluN2A subunit was also decreased in 5XFAD hippocampus (genotype effect, F_1,61_=6.31, p<0.05) (Figure 5D) and cortex (p<0.05) (Figure S4D) without change GluN2B and subunit ratios. These data demonstrate dysregulation to glutamate receptor expression beginning by 7 months of age in 5XFAD mice and an exacerbation of this effect due to PTZ kindling.

In addition, we used our retrospective quantitative measures of disease severity in this same cohort of mice^14^ and found that NKCC1/KCC2 was positively correlated with Aα_42_ (r=0.57, p<0.05) and pTau AT100 (Thr212, Ser214)/Tau (r=0.64, p<0.05) in the hippocampus of PTZ kindled 5XFAD mice and with performance on the Y-maze (r=-0.61, p<0.01) (Figure 5F, G, H) which we previously demonstrated was worsened by PTZ kindling. No correlations were found in non-kindled mice nor between GABA_A_R, AMPAR, or NMDAR in kindled mice. These findings suggest a relationship between inhibitory dysregulation, AD pathology, and cognitive decline.

When taken with the data from prodromal mice (Figure 3), 5XFAD mice show a worsening of markers of inhibitory imbalance, which was associated with worsened pathology and cognitive decline, and the emergence of dysregulated glutamate receptors as pathology progresses from prodrome to later stages. Further, PTZ kindling exacerbated the imbalance in both excitatory and inhibitory markers.

### mTORC1 inhibition modifies select seizure-induced excitation:inhibition markers in the hippocampus of 5XFAD mice

Rapamycin treatment has been shown to rescue neuronal hyperexcitability in various disease models ^53–58^. Thus, we sought to determine whether chronic low-dose rapamycin treatment beginning at the cessation of PTZ kindling (prodromal stage ∼4 months, 2.24 mg/kg daily), previously used to rescue seizure-induced neuropathology and cognitive deficits in 5XFAD mice^14^, could ameliorate PTZ-exacerbated E:I imbalance. The 5XFAD (Figure 6) and WT (Figure S5) mice used in our previous report were analyzed separately by two-way ANOVA with PTZ kindling and rapamycin treatment as independent variables.

**Figure 6.**
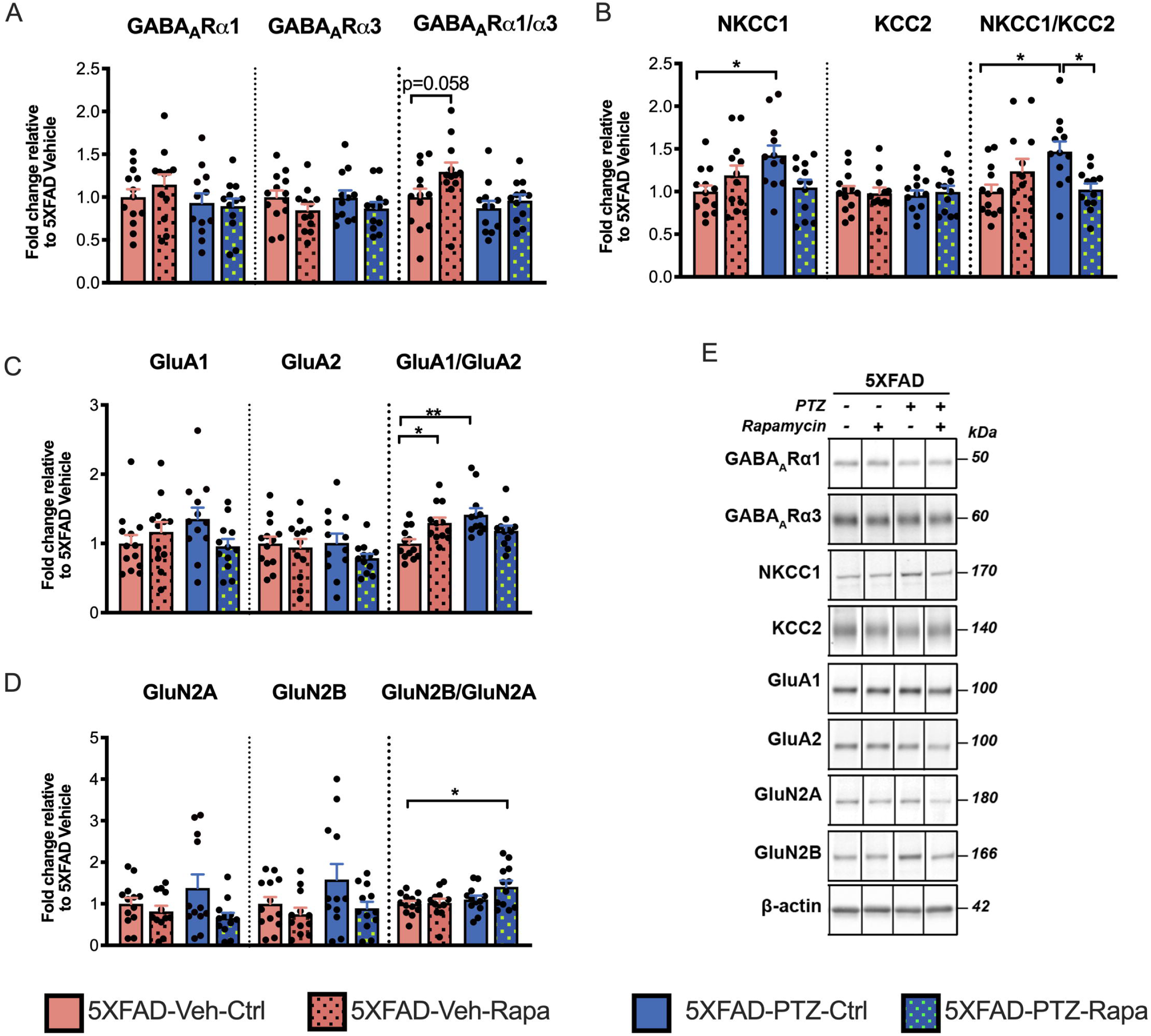
Differential effects of mTORC1 inhibition on seizure-induced excitation:inhibition imbalance in 5XFAD mice. (**A-D**) Quantification of (**A**) GABA_A_Rα1 and GABA_A_Rα3 and corresponding GABA_A_Rα1/GABA_A_Rα3 ratio, (**B**) Cl^-^ cotransporters NKCC1 and KCC2 and corresponding NKCC1/KCC2 ratio, (**C**) AMPAR subunits GluA1 and GluA2 and corresponding ratio GluA2/GluA1, and (**D**) NMDAR subunits GluN2A and GluN2B and corresponding GluN2B/GluN2A ratio. (**E**) Representative Western blot images for (**A-D**) showing non-adjacent bands originating from the same blot. n = 12-13 for each group. *p<0.05, **p<0.01.

Rapamycin showed a strong trend (p=0.058) to increase GABA_A_Rα1/α3 in non-kindled 5XFAD mice (kindling effect: F_1,45_=6.1, p<0.05; rapamycin effect: F_1,45_=4.1, p<0.05). These shifts corresponded with GABARα1 subunit expression (kindling effect: F_1,45_= 4.2, p<0.05; rapamycin effect: F_1,45_=6.1, p<0.05) (Figure 6A). Rapamycin also reversed the elevation of NKCC1/KCC2 ratio found in PTZ kindled 5XFAD mice (interaction: F_1,45_=9.8, p<0.01) compared to control 5XFAD mice (p<0.05) (Figure 6B). The changes in Cl^-^ cotransporter ratio was largely driven by changes in the Cl^-^ importer, NKCC1 (Figure 6B). These results likely reflect a rescue of GABA_A_R function in kindled 5XFAD mice by rapamycin.

The analysis of AMPAR subunits showed that PTZ increased GluA1/GluA2 ratios in 5XFAD mice (Figure 6C), consistent with the nontherapeutic cohort, and this was reversed by rapamycin treatment (Figure 6C). Rapamycin treatment increased GluA1/GluA2 ratio in non-kindled 5XFAD mice, largely driven by increased GluA1 expression, which reflected similar alterations (interaction: F_1,45_: F=4.3, p<0.05). Additionally, rapamycin treatment reduced GluN2A and GluN2B levels regardless of PTZ treatment (GluN2A rapamycin effect: F_1,45_=5.2, p<0.05; GluN2B rapamycin effect: F_1,45_=4.1, p<0.05) while PTZ kindled, rapamycin treated 5XFAD mice had elevated GluN2B/GluN2A ratio compared to control 5XFAD mice (Tukey’s post hoc: p<0.05) (Figure 6D). Overall, these data suggest that rapamycin can ameliorate PTZ-exacerbated inhibitory dysregulation in 5XFAD mice.

In WT mice, no significant differences were found due to PTZ kindling or rapamycin treatment for GABA_A_Rs, AMPARs, or NMDARs (Figure S5.), although kindling did increase NKCC1 expression in WT mice across rapamycin and vehicle treatments (Kindling effect: F_1,47_=5.2, p<0.05) (Figure S5B).

## Discussion

Here we identified novel mechanisms of neuronal dysfunction that may underlie seizure-exacerbated cognitive and functional deficits in AD. AD temporal cortex showed dysregulation of E:I balance proteins which was worsened in those with seizure history. Similarly, physiologic and protein indications of neuronal E:I imbalance were found in prodromal 5XFAD mice and further perturbations to synaptic protein regulators were seen at later stages and worsened in PTZ-kindled mice (Tables 1 and 2). Together with our recent study demonstrating enhanced AD pathology in AD+Sz patients and PTZ kindled mice^14^, our data suggest that E:I imbalance begins at prodromal stages and may play a role in seizure-induced exacerbation of neuropathology and cognitive decline in AD. Specific E:I alterations include decreased GABA_A_R subunit expression and shifts in the balance between the Cl^-^ cotransporters NKCC1/KCC2 and the AMPAR GluA1/GluA2 subunit ratios as factors in this interaction between AD and seizures (Tables 1 and 2). Importantly, we found that markers of E:I balance correlated with clinical measures of decline and disease severity in rare autopsy-confirmed tissue samples with prospective collection of seizure history data, justifying follow up studies to further examine clinical outcomes associated with seizure history in AD with more comprehensive statistical modeling that was not possible in the dataset here. Furthermore, our data indicate that using mTORC1 inhibitors such as rapamycin to therapeutically target the E:I imbalance may prove to be a promising therapy to alleviate AD progression due to neuronal hyperexcitability.

**Table 1.** Effects of AD and 5XFAD genotype on E:I balance. Prodromal = 4 months old.

|  |  | Prodromal 5XFAD | 7 month 5XFAD | Human |
| --- | --- | --- | --- | --- |
| GABA<sub>A</sub>R | Hippocampus | – GABA<sub>A</sub>R α1<br>– GABA<sub>A</sub>R α2<br>↓ GABA<sub>A</sub>R α1/α2 | ↓ GABA<sub>A</sub>R α1<br>– GABA<sub>A</sub>R α3<br>↓ GABA<sub>A</sub>R α1/α3 | ↓ GABA<sub>A</sub>R α1<br>– GABA<sub>A</sub>R α3<br>↓ GABA<sub>A</sub>R α1/α3 |
|  | Cortex | – GABA<sub>A</sub>R α1<br>– GABA<sub>A</sub>R α2<br>– GABA<sub>A</sub>R α1/α2 | ↓ GABA<sub>A</sub>R α1;<br>↓ GABA<sub>A</sub>R α3;<br>↓ GABA<sub>A</sub>R α1/α3 |  |
| Cl<sup>-</sup> cotransporters | Hippocampus | – NKCC1<br>– KCC2<br>– NKCC1/KCC2 | – NKCC1<br>– KCC2<br>– NKCC1/KCC2 | – NKCC1<br>– KCC2<br>– NKCC1/KCC2 |
|  | Cortex | – NKCC1<br>– KCC2<br>– NKCC1/KCC2 | ↓ NKCC1<br>– KCC2<br>– NKCC1/KCC2 |  |
| AMPA | Hippocampus | – GluA1<br>– GluA2<br>– GluA1/GluA2 | – GluA1<br>↓ GluA2<br>– GluA1/GluA2 | – GluA1<br>↑ GluA2<br>↓ GluA1/GluA2 |
|  | Cortex | – GluA1<br>– GluA2<br>– GluA1/GluA2 | – GluA1<br>↓ GluA2<br>– GluA1/GluA2 |  |
| NMDAR | Hippocampus | – GluN2A<br>– GluN2B<br>– GluN2B/GluN2A | ↓ GluN2A<br>– GluN2B<br>– GluN2B/GluN2A | ↑ GluN2A<br>↑ GluN2B<br>– GluN2B/GluN2A |
|  | Cortex | – GluN2A<br>– GluN2B<br>– GluN2B/GluN2A | ↓ GluN2A<br>– GluN2B<br>– GluN2B/GluN2A |  |

**Table 2.** Effects of seizures in AD and 5XFAD .mice.

|  |  | 7 month 5XFAD | Human AD |
| --- | --- | --- | --- |
| GABA<sub>A</sub>R | Hippocampus | —GABA<sub>A</sub>R α1<br>—GABA<sub>A</sub>R α 3<br>—GABA<sub>A</sub>R α1/α3 | ↓GABA<sub>A</sub>R α1<br>↓GABA<sub>A</sub>R α 3<br>—GABA<sub>A</sub>R α1/α3 |
|  | Cortex | —GABA<sub>A</sub>R α1<br>—GABA<sub>A</sub>R α 3<br>—GABA<sub>A</sub>R α1/α3 |  |
| Cl<sup>-</sup> cotransporters | Hippocampus | ↑NKCC1<br>—KCC2<br>↑NKCC1/KCC2 | —NKCC1<br>↓KCC2<br>↑NKCC1/KCC2 |
|  | Cortex | —NKCC1<br>—KCC2<br>—NKCC1/KCC2 |  |
| AMPA | Hippocampus | — GluA1<br>— GluA2<br>↑GluA1/GluA2 | —GluA1<br>—GluA2<br>—GluA1/GluA2 |
|  | Cortex | — GluA1<br>— GluA2<br>↑GluA1/GluA2 |  |
| NMDAR | Hippocampus | —GluN2A<br>— GluN2B<br>—GluN2B/GluN2A | —GluN2A<br>— GluN2B<br>—GluN2B/GluN2A |
|  | Cortex | —GluN2A<br>— GluN2B<br>—GluN2B/GluN2A |  |

Previous studies have found profound alterations in synaptic GABAergic signaling in AD including downregulation of GABA_A_R subunits^36,37,77,78^, decreased expression of inhibitory synaptic and perisomatic terminals^79 80^, indicative of reduced GABA_A_R synapses overall. Here we found decreased GABA_A_Rα1/α3 ratios in AD patients as compared to controls, supporting previous reports of reduced GABA sensitivity in membranes isolated from temporal cortex of AD brains^37^. In addition, AD patients with seizure history expressed significantly lower GABA_A_Rα1 and GABA_A_Rα3 subunits with GABA_A_Rα1 level correlating with brain atrophy in AD patients with seizures. NKCC1/KCC2 ratios were increased in AD with a history of seizures as compared to those without, consistent with collapse of neuronal Cl^-^gradient. In addition, we found a loss of PV containing interneurons in the temporal cortex of AD patients compared to controls, consistent with the previously reported reduction of PV cells within the AD parahippocampal gyrus^81^, which was driven by the AD+Sz group. Fast-spiking PV interneurons are major contributors to generation of gamma oscillations, which are disrupted in AD patients^82,83^. Greater loss of this GABAergic population in AD patients with epilepsy may promote network hyperexcitability and contribute to worsened cognitive and functional performance found in these subjects (Figure 1). Taken in the context of prior studies, these data suggest that a history of seizures in AD is associated with diminished GABAergic transmission via reduced GABA_A_R synapses, loss of interneurons, and possibly, depolarizing GABA response at those synapses that remain due to elevated intracellular Cl^-^.

Electrophysiological recordings in AD models, and 5XFAD mice in particular, are sparse, especially at stages in which plaque pathology has developed. Our electrophysiological data from the CA1 of prodromal 5XFAD mice demonstrate profound alterations in inhibitory synaptic transmission seen by reductions in mIPSC amplitude associated with increased neuronal excitability. Consistent with alterations in mIPSCs, we found decreased GABA_A_Rα1/ GABA_A_Rα2 ratios in the hippocampus of prodromal 5XFAD mice. Indeed reductions in GABARα1 and α5 transcript were found in patients with mild cognitive impairment suggesting GABA_A_R vulnerability at initial disease stages^84^.

Non-kindled 7 month old 5XFAD mice also showed decreased expression of GABA_A_Rα1, which we and others previously showed to be associated with decreased PV immunoreactivity^14,85–88^, suggesting a broader deficit of inhibition and reflecting our results in AD temporal cortex. Indeed, GABA_A_R loss has been linked to seizures and cognitive impairment in mouse models of AD^89–91^. This mechanism may be responsible for such changes in the 5XFAD model and underlie the correlation we found between the loss of GABA_A_Rα1 and increased atrophy in AD patients with seizures. Notably, we found that female sex was a predictor of lower GABA_A_Rα1/α3 ratios, which may be a co-contributor to worsened cognitive outcomes in females^92^. Our data also demonstrate that in 5XFAD mice, induced seizures can exaggerate GABAergic dysfunction increased NKCC1/KCC2 ratios, recapitulating our human data, which were reversed in rapamycin treated mice. Furthermore, our correlations establish a direct link between E:I balance and levels of AD pathology, supporting previous literature positing links between neuronal excitability and AD progression^93^.

With respect to glutamatergic neurotransmission, studies of AD hippocampus have highlighted decreased expression of GluA2 compared to controls^94–96^, a configuration known to increase AMPAR Ca^2+^ permeability^26,27,29,97,98^. Notably, the human AD temporal cortex we examined was late stage (Table S1) and we instead found an increase in GluA2 with a decrease in corresponding GluA1/GluA2 ratio in AD compared to controls (Figure 1E). These changes may represent a compensatory mechanism to neuronal hyperexcitability, but the significant GluA2 elevation may actually result in further impairments, given that Ca^2+^-permeable AMPAR are critical for the induction of late-phase LTP, essential for the formation of long-term memory^99,100^, as seen in neurodevelopmental disorder models^101^. We also found an increase in NMDAR subunits in AD patients, which is consistent with studies of the AD parietal cortex demonstrating increased excitatory to inhibitory synaptic ratio^102^. Elevated GluN2A in AD patients may again represent a compensatory mechanism, as lower GluN2A was associated with worsened CDR-SOB scores across all AD patients in our dataset, consistent with correlations in working memory performance and GluN2A in aged rats^103^. Elevated GluN2B is associated with epileptogenesis^15,29^ and extrasynaptic excitotoxic signaling in AD^104^. Further, elevations in AMPAR and NMDAR subunits are suggestive of elevated glutamatergic synapses overall, which may further tip E:I balance towards excitation.

While changes in AMPAR and NMDAR were not found in prodromal 5XFAD mice, we found elevated intrinsic excitability of CA1 neurons, which are the primary output area of hippocampus, together with decreased GABAergic signals suggest increased excitability of the hippocampal network. At later stages, we found increased expression of GluA1/GluA2 AMPAR subunit which was exacerbated by PTZ kindling and ameliorated with rapamycin treatment. Overall, these results suggest that AMPAR and NMDAR subunit composition is dynamic through AD progression and certain changes could be compensatory or neuroprotective from excitotoxicity, as often seen in acute epilepsy models^105,106^.

Recent epidemiologic research has underscored strong associations between epilepsy, late onset seizures, and AD^1,107–111^. In the past, seizures were assumed to be an unfortunate biproduct of AD. We and others have provided increasing preclinical and clinical evidence that seizures significantly contribute to neuropathology and cognitive decline and may also be a treatable component of this complex disease. Indeed, in a recent phase 2a clinical trial, levetiracetam, an antiepileptic drug, was shown to improve memory and executive function in AD patients with epileptiform activity^112^. Beyond seizure suppression, our data suggests that targeting specific E:I imbalances also holds therapeutic potential. Notably, a recent study has identified bumetanide, a NKCC1 antagonist, as a top candidate to treat AD, which when applied to AD model mice ameliorated transcriptomic, electrophysiological, and cognitive deficits^113^. Our data demonstrated significant exacerbation of NKCC1/KCC2 in both AD+Sz and seizure kindled mice. NKCC1/KCC2 ratios were also negatively correlated with performance related to spatial working memory in kindled 5XFAD mice, supporting the notion that restoration of the Cl^-^ gradient, perhaps with bumetanide, is therapeutically relevant, particularly for AD patients with seizure history.

We recently demonstrated that seizures exacerbate mTORC1 in AD patients and 5XFAD mice, and that chronic low dose rapamycin treatment is sufficient to ameliorate AD pathology and cognitive dysfunction in PTZ kindled 5XFAD mice^14^. Rapamycin is FDA approved and has been proven to benefit both AD and epilepsy models^14,114,115^. While rapalogs may affect numerous signaling pathways downstream of mTORC1 to attenuate disease course, including autophagy and neuroinflammation^116,117^, growing evidence demonstrates efficacy in regulating neuronal hyperexcitability^53–58^. In the studies presented here, rapamycin reduced NKCC1/KCC2 and GluA1/GluA2 ratios in PTZ kindled 5XFAD mice and showed a trend to increase GABA_A_Rα1/GABA_A_Rα3 in non-kindled 5XFAD mice, suggesting reversal of seizure-exacerbated E:I imbalance, which may indicate restoration of proper circuitry involved in memory in these mice^14^. However, the chronic rapamycin treatment induced small, but significant elevations in GluN2B/GluN2A in PTZ kindled 5XFAD mice and increased GluA1/GluA2 in non-kindled 5XFAD mice. In contrast, studies that used acute rapamycin treatments demonstrated that mTORC1 blockade decreases GluN2B and GluA1^118–121^. Thus, the relative increases in GluN2B and GluA1 that we found may represent a compensatory mechanism in response to chronic administration in 5XFAD mice. Indeed, one prior study demonstrated that chronic rapamycin induced increases in surface GluN2B, and that these changes were associated with amelioration of age-dependent cognitive decline^122^. These data indicate that targeting E:I balance with rapamycin, should be explored for clinical efficacy in AD patients with seizure history.

In summary, the data presented here identified novel mechanisms of neuronal dysfunction in AD that are amplified by seizures and may play a role in worsened cognitive outcomes associated with seizures in AD. These data suggest that targeting E:I imbalance, perhaps with rapamycin as well as other antiepileptic agents, holds therapeutic promise and is particularly relevant for AD patients with comorbid seizures.

## Acknowledgements

The authors thank the CNDR at the University of Pennsylvania for supplying the human tissue utilized in these studies as well as the University of Maryland Brain and Tissue Bank, part of the NIH Neuro Biobank for additional control brain samples. We would also like to thank Dr. Robert Vassar for supplying some of the 4-month 5XFAD mouse tissue.

## Funding

These studies were supported by the National Institutes of Health (NIH) National Institute of Neurological Disorders and Stroke (NINDS): R21NS105437 (FEJ), R01NS101156 (DMT), National Institute on Aging (NIA): R01AG077692 (FEJ & DMT), and Institutional National Research Service Award T32AG000255 (AJB), Alzheimer’s Association AARF-22-972333 (AJB). Clinical and postmortem tissue and data from Alzheimer’s disease patients was obtained from NIH studies at the University of Pennsylvania including P30-AG-072979 and R01-AG-054519.

## Competing interests

The authors declare no conflicts of interest.

## Authors’ contributions

AJB, SG, DMT, FEJ conceived the studies in this manuscript. AJB, SG, XL, DS acquired and analyzed the mouse and human tissue data. AJB, DJI designed the human clinical analysis. EL performed and analyzed electrophysiological experiments. AJB, DJI, FEJ, DMT interpreted the data from mouse and human studies. AJB, DMT, FEJ wrote the manuscript. All authors read and approved the final manuscript.

## Supplementary material

Supplementary material can be found in a separate PDF file.

## Notes

### Competing Interest Statement

The authors have declared no competing interest.

### Summary of Updates

Additional experiments were added to the manuscript along with adjustments to text.

