## Supplemental Materials for "Excitatory: inhibitory imbalance in Alzheimer’s disease is exacerbated by seizures with attenuation after rapamycin treatment in 5XFAD mice"

### Supplementary material

### Materials and Methods

#### Neurocognitive assessment of AD patients

The global CDR was calculated based on testing of six cognitive domains on a 0-3 scale with a score of 0 indicating cognitively normal and 3 indicating severe dementia[1]. The CDR-SOB represents the sum score of the six domains (memory, orientation, judgement and problem solving, community affairs, home and hobbies, and personal care), with scores ranging from 0 to 18, which offers a more detailed measure of cognitive and functional impairment[2]. In addition, the maximum score on the Dementia Severity Rating scale (DSRS)[3] was retrieved for each subject in our banked tissue set. The DSRS is a questionnaire completed by the patient's caretaker to rate the patient's memory (0-6), recognition of family members (0-5), orientation to place (0-4), social and community activity (0-5), personal care (0-3), speech and language (0-6), orientation to time (0-4), ability to make decisions (0-4), home activities and responsibilities (0-4), eating (0-3), and mobility (0-6) with higher scores indicating more severe deficits. Scores from each domain are totaled to determine overall functional performance. No significant differences between AD-Sz and AD+Sz were found in years since onset to final CDR assessment (mean AD-Sz= 8.3 years; mean AD+Sz=9.9 years).

#### Y-maze spontaneous alternation test

The Y maze is used to measure short term memory given the willingness of mice to explore the novel arm of the maze. Mice underwent behavioral assessment at 6.5 months of age [4]. Mice were allowed to explore the Y-shaped maze with three arms of equal length (38.1 cm x 7.6 cm x 12.7 cm) and were recorded for 8 minutes. Spontaneous alternation was calculated as the number of arm entries/(total number of arms entered-2 ) x 100.

#### Immunohistochemistry

Human tissue was formalin fixed, paraffin embedded, and sectioned at 6 μm. Immunohistochemistry was performed using previously established protocols [4,5] with anti-PV antibodies (Table S2). Briefly, slides were incubated in primary antibody for 15 min and IgG and

HRP-linker conjugates for 8 min. Slides were then exposed to hydrogen peroxide for 5 min before being incubated with 3,3'-diaminobenzidine tetrahydrochloride hydrate (DAB) for 10 min. Finally, slides were counterstained for nuclei with hematoxylin before being mounted with permanent medium. Slides were then imaged with a Nikon Eclipse 80i microscope with a digital Nikon DS-Fi2 camera (Micro Video Instruments, Avon, MA) at 20x objective in a zigzag sequence to sample up to 14 non-overlapping images of the grey matter.

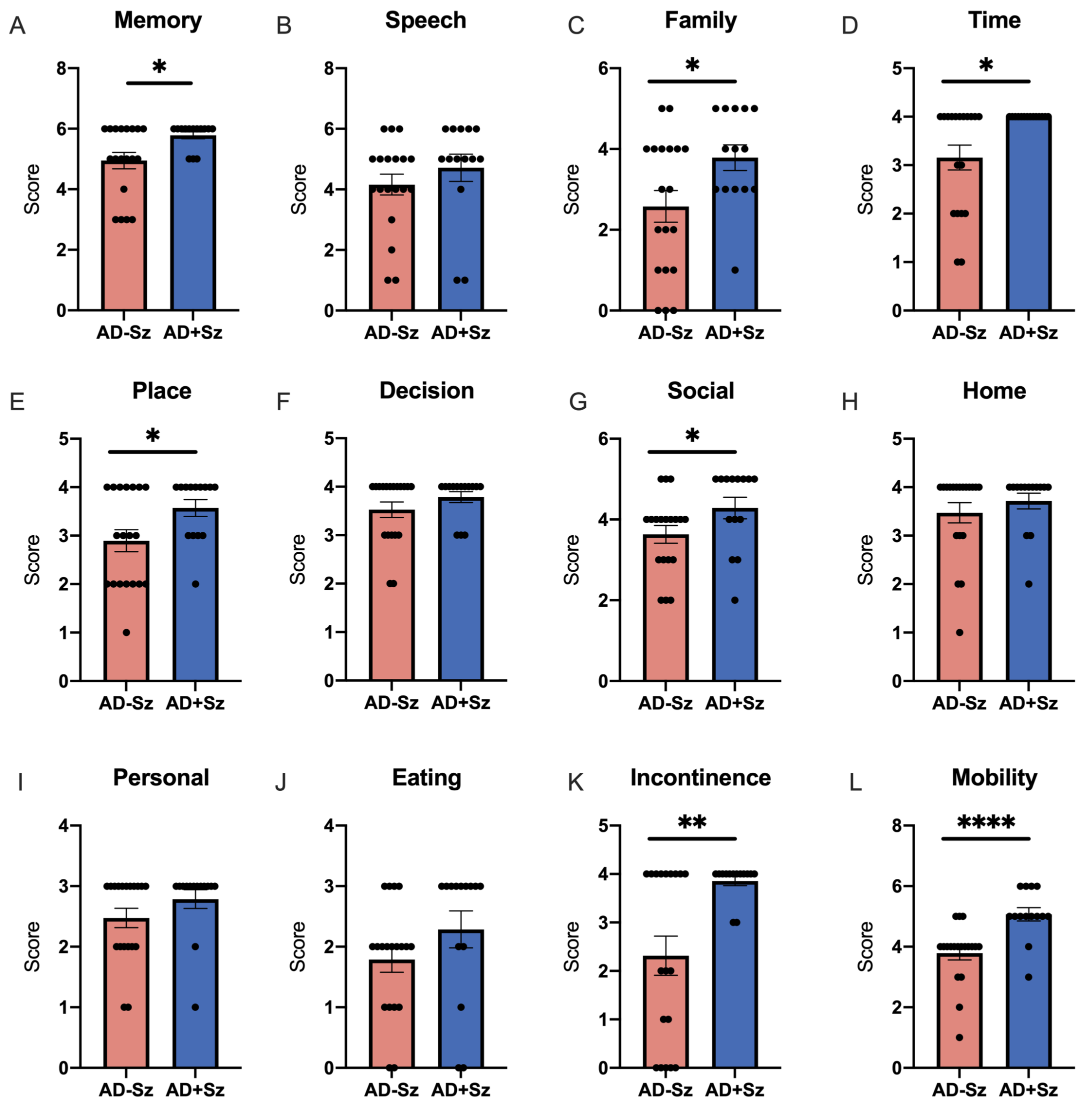


**Figure S1.** AD+Sz show increased deficits across several domains of the DSRS. Caretaker's ratings of patient's performance in (**A**) memory, (**B**) speech, (**C**) recognition of family, (**D**) orientation to time, (**E**) orientation to place, (**F**) decision making, (**G**) social and community activity, (**H**) home activities and responsibilities, (**I**) personal care, (**J**) eating, (**K**) incontinence, and (**L**) mobility. n=14-19, *p<0.05, **p<0.01, ****p<0.0001.

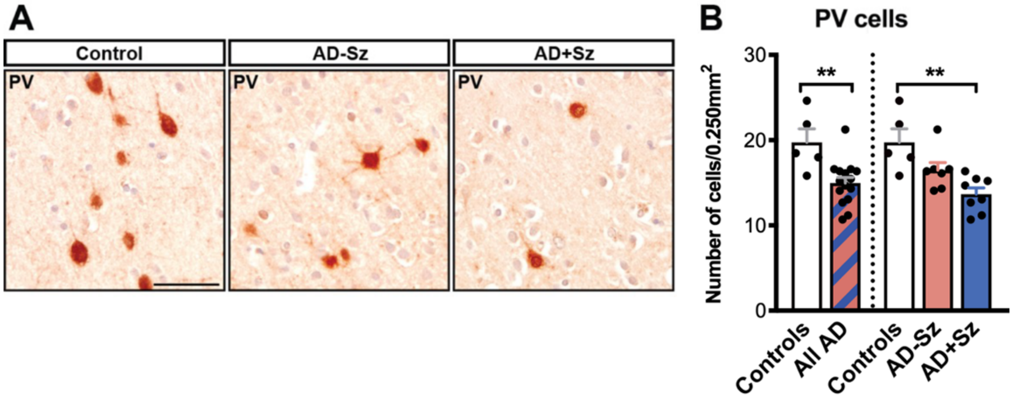


**Figure S2. PV cell loss in AD patients.** (**A**) Representative images of the temporal cortex from a control subject, an Alzheimer's disease (AD) patient without seizure history (AD-Sz) and an AD case with confirmed seizure history (AD+Sz), immunohistochemically labeled with parvalbumin (PV) showing an apparent decreased number of PV+ cells in AD cases compared to controls and further decreases in the AD+Sz compared to AD-Sz case. Scale bar represents 50 µm. (**B**) Number of PV positive cells per 0.250 $mm^2$ area counted in the temporal cortex of controls (n=5), all AD (n=15), and AD-Sz (n=7) and AD+Sz (n=8) subgroups. Control group is compared to all AD patients using t-test or compared to AD-Sz and AD+Sz groups using a one-way ANOVA. **$p<0.01$.

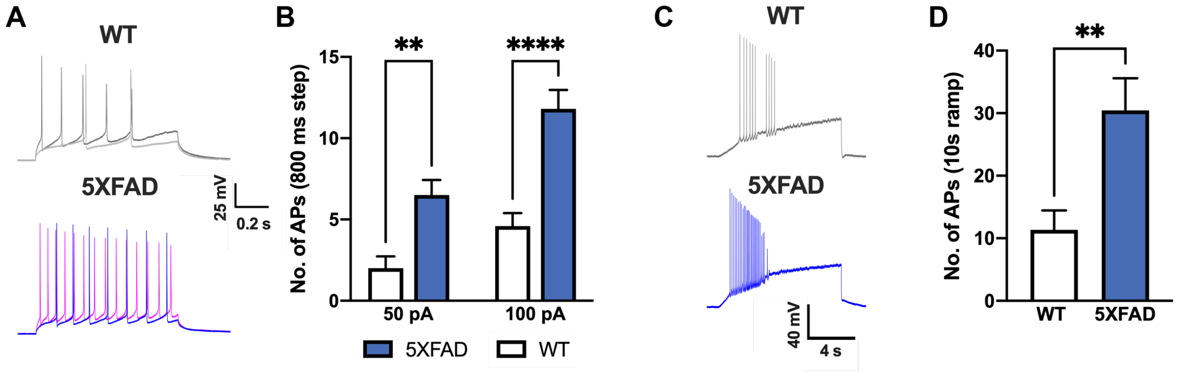


**Figure S3. Increased intrinsic excitability in pyramidal neurons from prodromal 5XFAD CA1.** (**A**) Representative traces and (**B**) quantification showing the number of APs evoked was significantly increased in 5XFAD pyramidal CA1 neurons compared to WT at both 50 pA (light grey, blue) and 100 pA (dark grey, red) current injection. n= 15 cells from 3 WT mice, 18 cells from 3 5XFAD mice. (**C**) Representative traces and (**D**) quantifications showing increased AP firing from ramp current injection in 5XFAD compared to WT pyramidal CA1 neurons. n= 15 cells from 3 WT mice, 17 cells from 3 5XFAD mice. **p<0.01, ****p<0.0001.

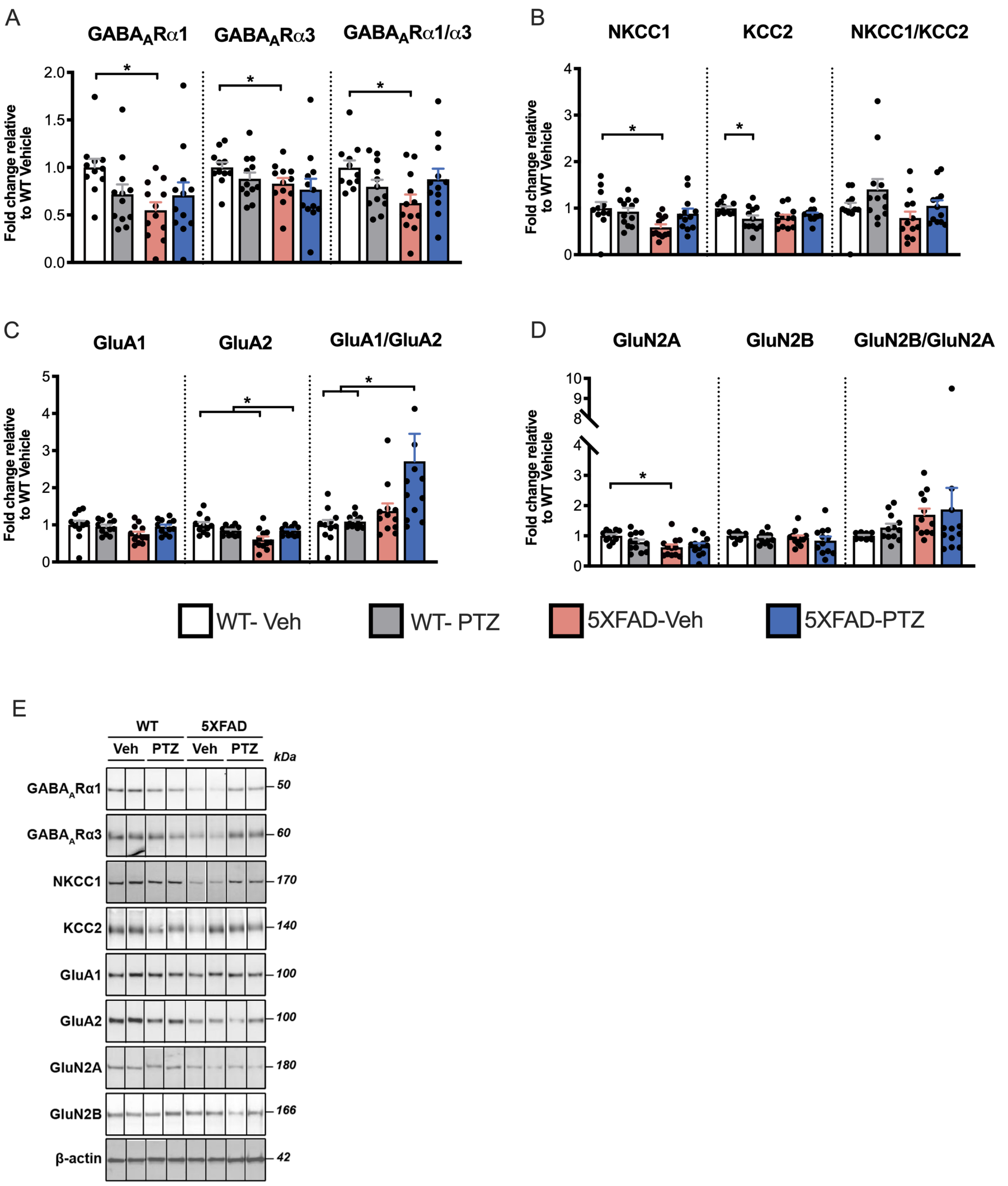


**Figure S4. Excitatory: inhibitory imbalance in the cortex of 5XFAD mice following induced seizures.** (**A-E**) Quantification of (**A**) GABA$_A$R subunits GABA$_A$Rα1 and GABA$_A$Rα3 and corresponding ratio GABA$_A$Rα1/GABA$_A$Rα3; (**B**) Cl$^-$ cotransporters NKCC1 and KCC2 and corresponding ratio NKCC1/KCC2; (**C**) AMPAR subunits GluA1 and GluA2 and corresponding ratio GluA2/GluA1; (**D**) NMDAR subunits GluN2A and GluN2B and corresponding ratio GluN2B/GluN2A. n = 12-19 WT-vehicle, 12-17 WT-PTZ, 12 5XFAD-vehicle and 17 5XFAD-PTZ. $^{*}p<0.05$, $^{**}p<0.01$.

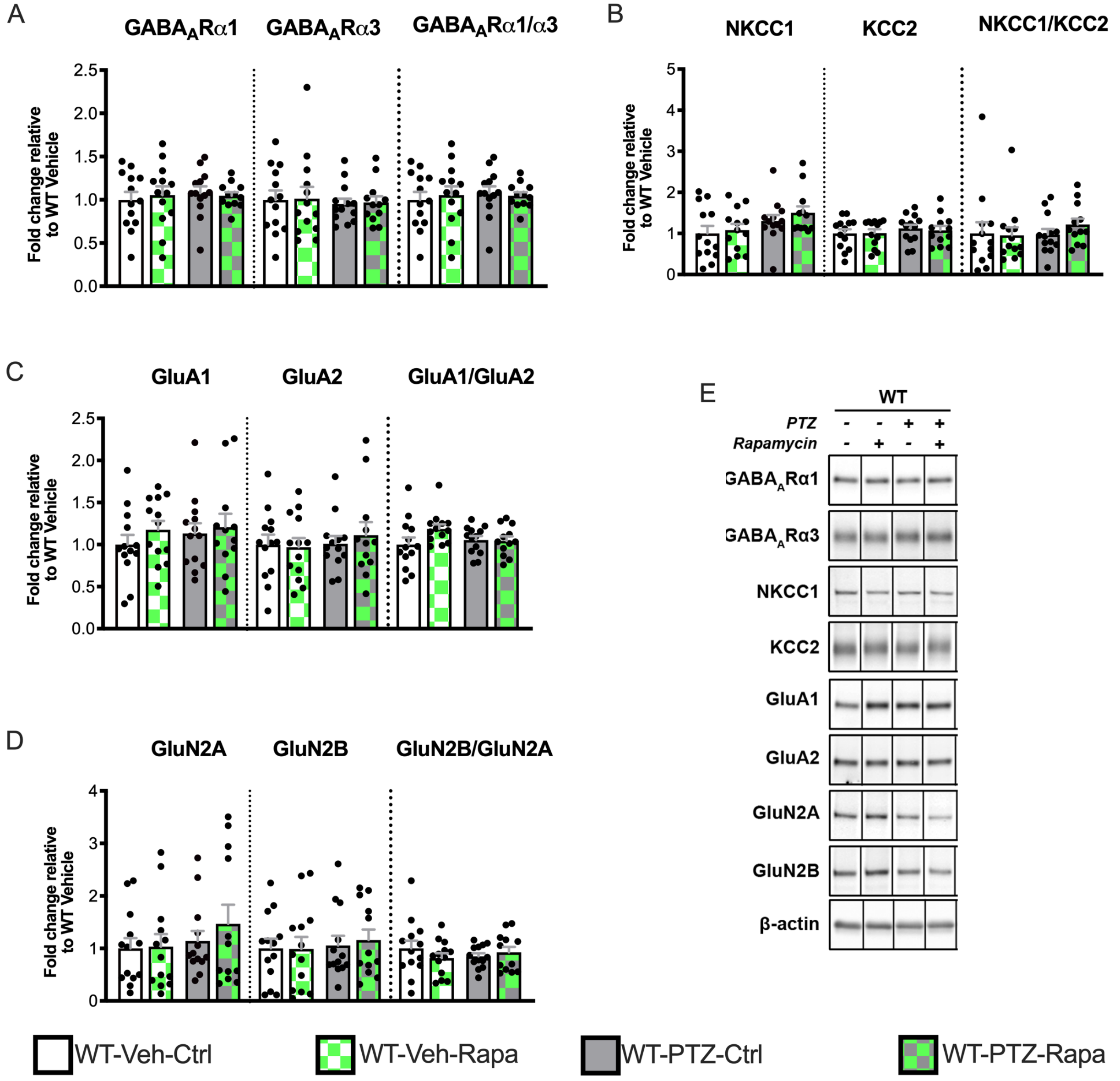


**Figure S5. E:I balance in control and rapamycin treated WT mice.** (**A-E**) Quantification of (**A**) $GABA_AR\alpha1$ and $GABA_AR\alpha3$ and corresponding $GABA_AR\alpha1/GABA_AR\alpha3$ ratio, (**B**) $Cl^-$ cotransporters NKCC1 and KCC2 and corresponding NKCC1/KCC2 ratio, (**C**) AMPAR subunits GluA1 and GluA2 and corresponding ratio GluA2/GluA1, and (**D**) NMDAR subunits GluN2A and GluN2B and corresponding GluN2B/GluN2A ratio. (**E**) Representative western blot images for (A-D) showing non-adjacent bands originating from the same blot. n = 12-13 for each group.

| **Pateint** | **Sex** | **Age at death (yr)** | **Disease duration (yr)** | **PMI (h)** | **Seizure history** | **Brain weight (g)** | **Thal stage** | **Braak stage** | **CERAD** |
|---|---|---|---|---|---|---|---|---|---|
| Ctrl1* | F | 46 | NA | 12 | unk | 1228 | 0 | 0 | NA |
| Ctrl2* | M | 57 | NA | 24 | unk | 1297 | 0 | 0 | 3 |
| *Ctrl3** | M | 57 | NA | 14 | unk | 1360 | 1 | 0 | NA |
| Ctrl4* | M | 59 | NA | 17 | unk | 1218 | 0 | 0 | NA |
| Ctrl5* | M | 59 | NA | 18 | unk | 1502 | 0 | 0 | NA |
| Ctrl6* | M | 62 | NA | 8.5 | unk | 1420 | 0 | 0 | 3 |
| *Ctrl7** | F | 65 | NA | 19 | unk | 1207 | 1 | 0 | 3 |
| *Ctrl8** | M | 67 | NA | 15 | unk | 1545 | 1 | 1 | NA |
| Ctrl9* | M | 68 | NA | 17 | unk | 1371 | 0 | 0 | 3 |
| Ctrl10* | M | 68 | NA | 14 | unk | 1512 | 0 | 0 | 3 |
| Ctrl11* | M | 70 | NA | 12 | unk | NA | NA | NA | NA |
| Ctrl12* | M | 72 | NA | 17 | unk | 1216 | 0 | 1 | NA |
| Ctrl13* | M | 75 | NA | 17 | unk | NA | NA | NA | 3 |
| *Ctrl 14* | F | 60 | NA | 16 | unk | 1374 | NA | NA | NA |
| *Ctrl 15* | F | 51 | NA | 16 | unk | 1143 | NA | NA | NA |
| **AD-Sz1** | F | 84 | 6 | 24 | unk | 1204 | NA | 6 | 3 |
| **AD-Sz2** | M | 90 | 14 | 19 | unk | 1309 | 4 | 6 | 3 |
| **AD-Sz3** | M | 78 | 8 | 6 | unk | 1190 | NA | 6 | 3 |
| **AD-Sz4** | M | 70 | 9 | 5 | unk | 1256 | NA | 6 | 3 |
| **AD-Sz5** | F | 90 | 20 | 84 | unk | 908 | NA | 6 | 3 |
| **AD-Sz6** | M | 78 | 11 | 3 | unk | 1199 | 5 | 5 | 3 |
| ***AD-Sz7**** | M | 79 | 7 | 17 | unk | 1345 | 4 | 6 | 3 |
| **AD-Sz8** | F | 88 | 12 | 20 | unk | 1094 | NA | NA | 3 |
| **AD-Sz9** | M | 87 | 14 | 19 | unk | 1198 | 5 | 6 | 3 |
| **AD-Sz10** | M | 86 | 14 | 10 | unk | 1278 | 5 | 6 | 3 |
| **AD-Sz11** | F | 62 | 6 | 8 | unk | 988 | NA | 6 | 3 |
| **AD-Sz12** | F | 90 | 7 | 17 | unk | 1124 | NA | NA | 3 |
| **AD-Sz13** | F | 79 | 10 | 4 | unk | 909 | 5 | 6 | 3 |
| **AD-Sz14** | M | 83 | 14 | 4 | unk | 1155 | 4 | 6 | 3 |
| **AD-Sz15** | F | 85 | 9 | 9 | unk | 1014 | 5 | 5 | 3 |
| **AD-Sz16** | M | 84 | 12 | 21 | unk | 1265 | 2 | 3 | 3 |
| **AD-Sz17** | M | 44 | 2 | 44 | unk | 1510 | NA | 6 | 3 |
| **AD-Sz18** | F | 85 | 7 | 14 | unk | 1138 | NA | 5 | 3 |

| | | | | | | | | | |
|---|---|---|---|---|---|---|---|---|---|
| **AD-Sz19*** | M | 63 | 9 | 16 | unk | 1243 | NA | 6 | 3 |
| **AD-Sz20** | F | 74 | 12 | 4 | unk | 989 | 5 | 6 | 3 |
| **AD-Sz21** | M | 98 | 10 | 48 | unk | 1438 | NA | 3 | 3 |
| **AD-Sz22** | F | 84 | 15 | 21.5 | unk | 1151 | 5 | 6 | 3 |
| **AD-Sz23** | M | 70 | 11 | 11 | unk | 1082 | NA | 6 | 3 |
| **AD-Sz24*** | F | 70 | 13 | 6 | unk | 978 | NA | NA | 3 |
| **AD-Sz25*** | M | 71 | 12 | 4 | unk | 1330 | NA | 6 | 3 |
| **AD-Sz26** | M | 84 | 7 | 18 | unk | 1281 | NA | 5 | 3 |
| **AD-Sz27** | F | 85 | 5 | 20 | unk | 1026 | NA | NA | 3 |
| **AD-Sz28** | M | 78 | 10 | 18 | unk | 1100 | NA | 6 | 3 |
| ***AD-Sz29**** | F | 65 | 10 | 8.5 | unk | 997 | 5 | 6 | 3 |
| **AD-Sz30** | M | 84 | 16 | 18.5 | unk | 1185 | 5 | 6 | 3 |
| **AD-Sz31** | F | 98 | 12 | 24 | unk | 938 | NA | 6 | 3 |
| **AD-Sz32** | F | 90 | 11 | 12 | unk | 1269 | 5 | 6 | 3 |
| **AD-Sz33** | F | 74 | 10 | 6 | unk | 1149 | 5 | 6 | 3 |
| **AD-Sz34** | F | 70 | 19 | 16 | unk | 845 | 5 | 6 | 3 |
| ***AD-Sz35**** | M | 64 | 9 | 16.5 | unk | 1320 | NA | 6 | 3 |
| **AD-Sz36** | M | 84 | 15 | 7 | unk | 1309 | NA | 6 | 3 |
| **AD-Sz37** | M | 78 | 14 | 6.5 | unk | 926 | NA | 6 | 3 |
| **AD-Sz38** | M | 65 | 6 | 17 | unk | 960 | NA | 6 | 3 |
| **AD-Sz39*** | F | 64 | 5 | 20 | unk | 1098 | NA | 3 | 2 |
| **AD-Sz40** | M | 72 | 9 | 8 | unk | 1167 | NA | 6 | 3 |
| **AD-Sz41** | M | 68 | 14 | 7 | unk | 1058 | NA | 6 | 3 |
| **AD-Sz42** | F | 88 | 10 | 15 | unk | 1145 | NA | 5 | 2 |
| **AD-Sz43** | F | 89 | 17 | 12 | unk | 1080 | NA | 6 | 3 |
| **AD-Sz44** | F | 91 | 15 | 12 | unk | 1283 | NA | 6 | 2 |
| **AD-Sz45** | F | 89 | 13 | 5 | unk | 1060 | NA | NA | 3 |
| **AD-Sz46** | F | 85 | 15 | 12 | unk | 1069 | 4 | 6 | 3 |
| **AD-Sz47** | M | 90 | 8 | 11 | unk | 1459 | 5 | 6 | 3 |
| ***AD-Sz48**** | M | 91 | 7 | 4 | unk | 1105 | NA | 6 | 3 |
| **AD-Sz49** | M | 75 | 14 | 18.5 | unk | 1210 | 5 | 6 | 3 |
| **AD-Sz50** | F | 84 | 10 | 5 | unk | 1053 | NA | NA | 3 |
| **AD-Sz51** | M | 81 | 8 | 18 | unk | 1427 | NA | 6 | 3 |
| **AD-Sz52** | M | 83 | 17 | 4.5 | unk | 1013 | 5 | 6 | 3 |
| **AD-Sz53*** | F | 84 | 6 | 11 | unk | 1043 | NA | 6 | 3 |
| **AD-Sz54** | M | 90 | 7 | 20 | unk | 1237 | NA | 5 | 3 |

| | | | | | | | | | |
|---|---|---|---|---|---|---|---|---|---|
| **AD-Sz55** | F | 64 | 11 | 14 | unk | 1099 | 5 | 6 | 3 |
| **AD-Sz56*** | M | 61 | 8 | 18 | unk | 1203 | 5 | 6 | 3 |
| **AD-Sz57** | M | 86 | 11 | 6.5 | unk | 1147 | 5 | 6 | 3 |
| **AD-Sz58** | M | 71 | 9 | 4.5 | unk | 1234 | 5 | 6 | 3 |
| **AD-Sz59** | F | 85 | 19 | 4 | unk | 969 | 5 | 6 | 2 |
| **AD-Sz60*** | F | 82 | 11 | 7 | unk | 1113 | 5 | 6 | 3 |
| ***AD-Sz61**** | F | 86 | 10 | 6 | unk | 889 | 5 | 5 | 2 |
| **AD-Sz62** | M | 85 | 3 | 6.5 | unk | 1301 | NA | 4 | 0 |
| **AD-Sz63** | M | 91 | 5 | 9 | unk | 1232 | 5 | 3 | 2 |
| **AD-Sz64** | M | 79 | 7 | 20 | unk | 1147 | NA | 6 | 3 |
| **AD-Sz65** | M | 85 | 20 | 16 | unk | 1404 | 5 | 5 | 3 |
| **AD-Sz66** | F | 83 | 17 | 11 | unk | 845 | NA | NA | 3 |
| **AD-Sz67** | F | 75 | 10 | 18 | unk | 1207 | 5 | 6 | 3 |
| **AD-Sz68** | M | 70 | 14 | 4 | unk | 1201 | NA | NA | 3 |
| ***AD-Sz69**** | F | 90 | 7 | 10 | unk | 1214 | NA | 6 | 3 |
| **AD-Sz70** | F | 85 | 24 | 4 | unk | 967 | NA | 6 | 3 |
| **AD-Sz71** | M | 75 | 7 | 13 | unk | 1403 | NA | 6 | 3 |
| **AD-Sz72** | F | 88 | 12 | 27 | unk | 1205 | NA | 5 | 1 |
| **AD-Sz73*** | F | 63 | 12 | 27 | unk | 999 | NA | NA | 3 |
| **AD-Sz74*** | M | 83 | 10 | 18 | unk | 1336 | 5 | 6 | 3 |
| **AD-Sz75** | F | 89 | 16 | 18 | unk | 1208 | NA | 6 | 3 |
| **AD-Sz76** | F | 88 | 16 | 7 | unk | 1016 | NA | NA | 3 |
| **AD-Sz77** | M | 88 | 9 | 17 | unk | 1177 | NA | 4 | 2 |
| **AD-Sz78** | F | 85 | 6 | 20 | unk | 1202 | 5 | 6 | 3 |
| **AD-Sz79** | F | 70 | 18 | 3.5 | unk | 1068 | NA | 6 | 3 |
| **AD-Sz80** | F | 86 | NA | 21 | unk | 1128 | NA | 6 | 3 |
| **AD-Sz81** | M | 75 | 10 | 19 | unk | 1193 | NA | 6 | 3 |
| **AD-Sz82** | F | 73 | 10 | 22 | unk | 1022 | 5 | 6 | 3 |
| **AD-Sz83** | F | 77 | 7 | 17 | unk | 1102 | 5 | 6 | 3 |
| **AD-Sz84** | M | 72 | 4 | 24 | unk | 1176 | 4 | 6 | 3 |
| **AD-Sz85** | F | 78 | NA | 12 | unk | 1116 | 5 | 5 | 3 |
| ***AD-Sz86**** | M | 79 | NA | 16 | unk | 1426 | 5 | 5/6 | 3 |
| **AD-Sz87*** | F | 72 | NA | 12 | unk | 1092 | 5 | 6 | 2 |
| AD-Sz88* | F | 63 | 10 | 5 | unk | 1137 | NA | NA | 3 |
| AD-Sz89* | M | 77 | 3 | 17 | unk | 1265 | NA | NA | 2 |
| ***AD+Sz1**** | M | 83 | 12 | 11 | Y | 1077 | NA | 6 | 3 |

| | | | | | | | | | |
|---|---|---|---|---|---|---|---|---|---|
| ***AD+Sz2**** | F | 68 | 8 | 9 | Y | 1000 | NA | 6 | 3 |
| **AD+Sz3*** | F | 80 | 13 | 12 | Y | 1063 | 5 | 6 | 2 |
| **AD+Sz4** | F | 68 | 12 | 12 | Y | 942 | 5 | 6 | 2 |
| **AD+Sz5*** | M | 71 | 8 | 20 | Y | 1164 | 5 | 6 | 3 |
| **AD+Sz6** | F | 83 | 24 | 7 | Y | 783 | 5 | 6 | 3 |
| **AD+Sz7** | F | 86 | 12 | 15 | Y | 1088 | NA | 4 | 0 |
| **AD+Sz8*** | F | 81 | 10 | 5 | Y | 847 | NA | 6 | 3 |
| **AD+Sz9*** | F | 77 | 6 | 3 | Y | 1010 | NA | 6 | 3 |
| **AD+Sz10** | F | 76 | 10 | 45 | Y | 1139 | 5 | 6 | 3 |
| ***AD+Sz11**** | M | 89 | 12 | 3 | Y | 1034 | NA | 6 | 3 |
| ***AD+Sz12**** | F | 65 | 10 | 11 | Y | 1125 | NA | 6 | 3 |
| **AD+Sz13** | M | 71 | 6 | 23 | Y | 1206 | NA | 6 | 3 |
| **AD+Sz14** | F | 77 | 10 | 10 | Y | 922 | NA | 6 | 3 |
| *AD+Sz15** | F | 76 | 18 | 11 | Y | 1125 | NA | 6 | NA |
| ***AD+Sz16**** | F | 85 | NA | 19 | Y | 953 | 4/5 | 5/6 | 2 |
| **AD+Sz17*** | F | 91 | 20 | 12 | Y | 980 | 5 | 6 | NA |
| *AD+Sz18** | M | 68 | NA | 5 | Y | 929 | NA | 5/6 | NA |
| **AD+Sz19*** | M | 79 | 17 | 8 | Y | 1005 | NA | 6 | NA |
| ***AD+Sz20**** | M | 84 | 15 | 19 | Y | 1150 | NA | 2 | NA |

**Table S1**. Human subject clinical information and neuropathology. Bold indicates subjects included in CDR, * indicates subjects included in western blot and DSRS, italics indicates subjects included in IHC. NA= data not available; unk = unknown.

| Antibody | Ref (Distributor) | Isotype | Size (kDa) |
|---|---|---|---|
| β-actin | A5441 (Sigma A.) | Mouse monoclonal IgG | 43 |
| Amyloid- β | 803001 (Biolegend) | Mouse monoclonal IgG1 | 4 |
| $GABA_AR$ α1 | AB5609 (Millipore) | Rabbit polyclonal IgG | 51 |
| $GABA_AR$ α2 | MABN1724 (Millipore) | Mouse monoclonal IgG1κ | 57 |
| $GABA_AR$ α3 | G4291 (Sigma A.) | Rabbit polyclonal IgG | 60 |
| GluA1 | ab31232 (abcam) | Rabbit polyclonal IgG | 100 |
| GluA2 | AB1768 (Millipore) | Rabbit polyclonal IgG | 108 |
| KCC2 | 07-432 (Millipore) | Rabbit polyclonal IgG | 140 |
| NKCC1 | AB3560P (Millipore) | Rabbit polyclonal | 170 |
| GluN2A | M264 (Sigma A.) | Rabbit polyclonal | 180 |
| GluN2B | MA1-2014 (Thermo Sci.) | Mouse monoclonal IgG1 | 166 |
| PV | PA1-933 (Thermo Sci.) | Rabbit polyclonal IgG | 12 |
| Tau 5 | AHB0042 (Thermo Sci.) | Mouse monoclonal IgG1 | 50 |
| p-Tau AT100 | MN1060 (Thermo Sci.) | Mouse monoclonal IgG1 | 40-80 |

**Table S2**. Primary antibody information.
